# Beyond Drift Diffusion Models: Fitting a broad class of decision and RL models with HDDM

**DOI:** 10.1101/2022.06.19.496747

**Authors:** Alexander Fengler, Krishn Bera, Mads L. Pedersen, Michael J. Frank

## Abstract

Computational modeling has become a central aspect of research in the cognitive neurosciences. As the field matures, it is increasingly important to move beyond standard models to quantitatively assess models with richer dynamics that may better reflect underlying cognitive and neural processes. For example, sequential sampling models (SSMs) are a general class of models of decision making intended to capture processes jointly giving rise to reaction time distributions and choice data in n-alternative choice paradigms. A number of model variations are of theoretical interest, but empirical data analysis has historically been tied to a small subset for which likelihood functions are analytically tractable. Advances in methods designed for likelihood-free inference have recently made it computationally feasible to consider a much larger spectrum of sequential sampling models. In addition, recent work has motivated the combination of SSMs with reinforcement learning (RL) models, which had historically been considered in separate literatures. Here we provide a significant addition to the widely used HDDM Python toolbox and include a tutorial for how users can easily fit and assess a (user extensible) wide variety of SSMs, and how they can be combined with RL models. The extension comes batteries included, including model visualization tools, posterior predictive checks, and ability to link trial-wise neural signals with model parameters via hierarchical Bayesian regression.

## 1 Introduction

The drift diffusion model (DDM, also called diffusion decision model or Ratcliff diffusion model) (Ratcliff, 1978; Ratcliff et al., 2016), and more generally the framework of sequential sampling models (SSMs) (Hawkins et al., 2015; Heathcote et al., 2022; Ratcliff et al., 2016; Tillman et al., 2020; Voss et al., 2019) have become a mainstay of the cognitive scientist’s model arsenal in the last two decades (Lawlor et al., 2020; Van Zandt et al., 2000; Wieschen et al., 2020).

SSMs are used to model neurocognitive processes that jointly give rise to choice and reaction time data in a multitude of domains, spanning from perceptual discrimination to memory retrieval to preference-based choice (Krajbich et al., 2012; Krajbich & Rangel, 2011; Ratcliff, 1978; Ratcliff et al., 2006; Smith et al., 2014) across species (Doi et al., 2020; Gold & Shadlen, 2007; Yartsev et al., 2018). Moreover, researchers are often interested in the underlying neural dynamics that give rise to such choice processes. As such, many studies include additional measurements such as EEG, fMRI or eyetracking signals as covariates, which act as latent variables and connect to model parameters (e.g. via a regression model) to drive trial specific parameter valuations (Doi et al., 2020; Forstmann et al., 2010; Frank et al., 2015; Rangel et al., 2008; Yartsev et al., 2018). See Figure 1 for an illustration of the DDM and some canonical experimental paradigms.

**Figure 1.**
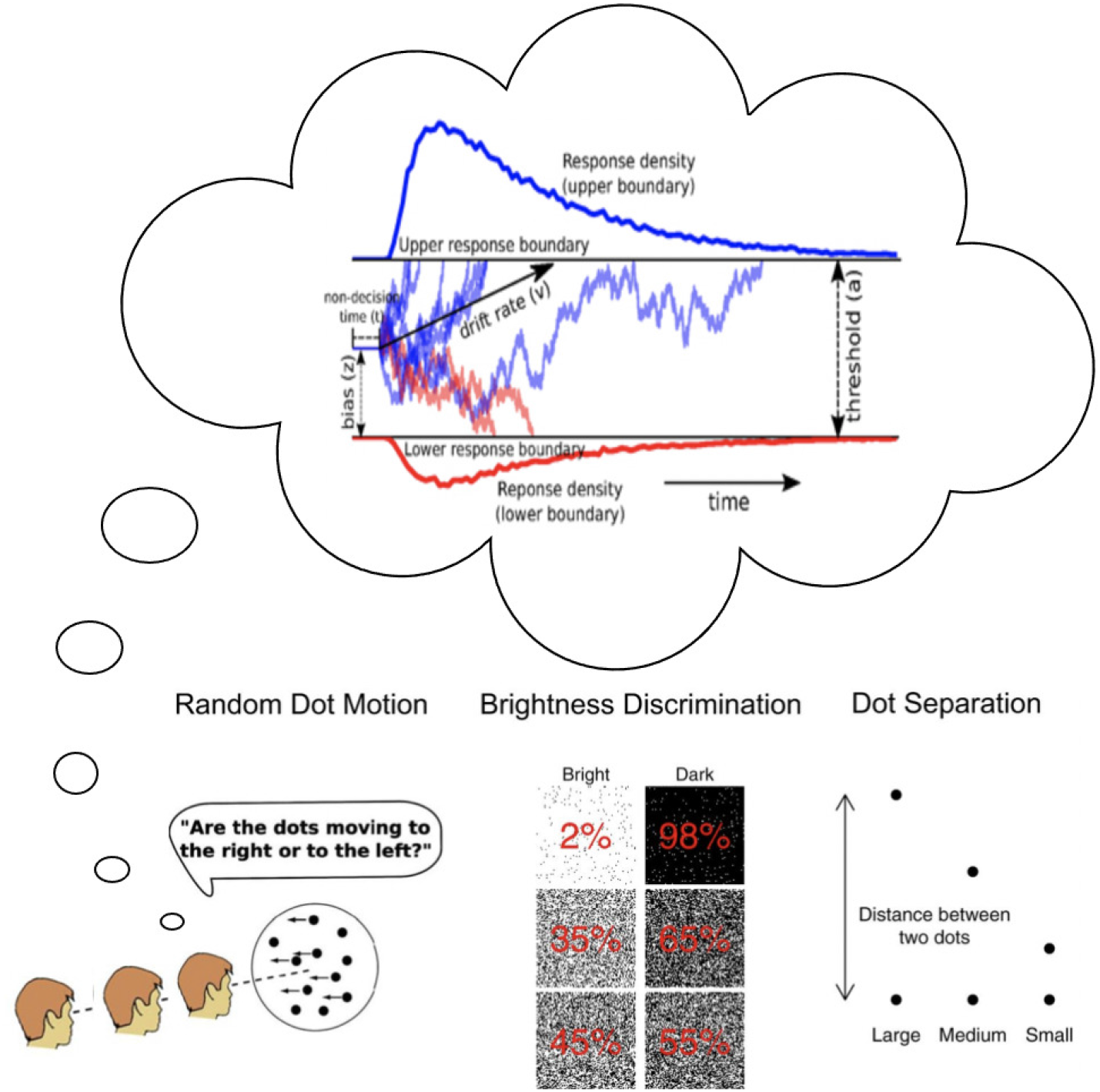
Drift Diffusion Model and some example applications.

The widespread interest and continuous use of SSMs across the research community has spurred the development of several software packages targeting the estimation of such models (Fontanesi, 2022; Heathcote et al., 2019; Vandekerckhove & Tuerlinckx, 2008). For a hierarchical Bayesian approach to parameter estimation, the HDDM toolbox in Python (Wiecki et al., 2013) (available at https://github.com/hddm-devs/hddm) is widely used and the backbone of hundreds of studies published in peer reviewed journals.

HDDM allows users to conveniently specify and estimate DDM parameters for a wide range of experimental designs, including the incorporation of trial-by-trial covariates via regression models targeting specific parameters. As an example, one may use this framework to estimate whether trial-by-trial drift rates in a DDM co-vary with neural activity in a given region (and/or temporal dynamic), pupil dilation or eye gaze position. Moreover, by using hierarchical Bayesian estimation, HDDM optimizes the inference about such parameters at the individual subject and group levels.

Nevertheless, until now, HDDM and other such toolboxes have been largely limited to fitting the 2-alternative choice DDM (albeit allowing for the full DDM with inter-trial parameter variability). The widespread interest in SSMs has however also spurred theoretical and empirical investigations into various alternative model variants. Notable examples are, amongst others, race models with more than 2 decision options, the leaky competing accumulator model (Usher & McClelland, 2001), SSMs with dynamic decision boundaries (Cisek et al., 2009; Ratcliff & Frank, 2012; Trueblood et al., 2021) and more recently SSMs based on Levy flights rather than basic Gaussian diffusions (Wieschen et al., 2020). Moreover, as mentioned earlier, SSMs naturally extend to n-choice paradigms.

A similar state of affairs is observed for another class of cognitive models which aim to simultaneously model the dynamics of a feedback-based learning across trials as well as the within-trial decision process. One way to achieve this is by replacing the choice rule in a reinforcement learning (RL) process, in itself an important theoretical framework in the study of learning behavior across trials (Collins & Shenhav, 2022; Dowd et al., 2016; Eckstein et al., 2021; McDougle & Collins, 2021) with a cognitive process models such as SSMs. This forms a powerful combination of modeling frameworks. While recent studies moved into this direction (Fontanesi, 2022; Fontanesi et al., 2019; Pedersen & Frank, 2020; Pedersen et al., 2017; Turner, 2019), they have again been limited to an application of the basic DDM.

Despite the great interest in these classes of models, tractable inference and therefore, widespread adoption of such models has been hampered by the lack of easy to compute likelihood functions (including essentially all of the examples provided above). In particular, while many interesting models are straightforward to simulate, often researchers want to go the other way: from the observed data to infer the most likely parameters. For all but the simplest models, such likelihood functions are analytically intractable, and hence previous approaches required computationally costly simulations and/or lacked flexibility in applying such methods to different scenarios (Boehm et al., 2021; Palestro et al., 2019; Shinn et al., 2020; Turner & Sederberg, 2014; Turner & Van Zandt, 2018). We recently developed a novel approach using artificial neural networks which can, given sufficient training data, approximate likelihoods for a large class of SSM variants, thereby amortizing the cost and enabling rapid, efficient and flexible inference (Fengler et al., 2021). We dubbed such networks LANs, for *likelihood approximation networks*.

The core idea behind computation amortization is to run an expensive process only once, so that the fruits of this labor can later be reused and shared with the rest of the community. Profiting from the computational labor incurred in other research groups enables researchers to consider a larger bank of generative models and to sharpen conclusions that may be drawn from their experimental data. The benefit is three-fold. Experimenters will be able to adjudicate between a rising number of competing models (theoretical accounts), capture richer dynamics informed by neural activity, and at the same time new models proposed by theoreticians can find wider adoption and be tested against data much sooner.

Just as streamlining the analysis of simple SSMs (via e.g., the HDDM toolbox and others) allowed an increase in adoption, streamlining the production and inference pipeline for amortized likelihoods, we hope, will drive the embrace of SSM variations in the modeling and experimental community by making a much larger class of models ready to be fit to experimental data.

Here we develop an extension to the widely used HDDM toolbox, which generalizes it to allow for flexible simulation and estimation of a large class of SSMs by reusing amortized likelihood functions.

Specifically, this extension incorporates,

- LAN (Fengler et al., 2021) based likelihoods for a variety of SSMs (batteries included)
- LAN-driven extension of the Reinforcement Learning (RL) - DDM capabilities, which allows RL learning rules to be applied to all included SSMs
- New plots which focus on visual communications of results across models
- An easy interface for users to import and incorporate their own models and likelihoods into HDDM

This paper is formulated as a tutorial to support application of the HDDM LAN extension for data analysis problems involving SSMs.

The rest of the paper is organized as follows. In section 2 we start by providing some basic overview of the capabilities of HDDM. Section 3 gives a brief overview of LANs (Fengler et al., 2021). Section 4 constitutes a tutorial with a detailed introduction on how to use these new features in HDDM. We conclude in section 5 embedding the new features into a broader agenda. Lastly we mention limitations and preview future developments in section 6.

## 2 HDDM: The basics

The HDDM Python package (Wiecki et al., 2013) was designed to make hierarchical Bayesian inference for drift diffusion models simple for end-users with some programming experience in Python. The toolbox has been widely used for this purpose by the research community and the feature set evolves to accommodate new use-cases. This section serves as a minimal introduction to HDDM to render the present tutorial self-contained. To get a deeper introduction to HDDM itself, please refer to the original paper (Wiecki et al., 2013), an extension paper specifically concerning reinforcement learning capabilities (Pedersen & Frank, 2020), and the [documentation](https://hddm.readthedocs.io/en/latest/) of the package. Here we concern ourselves with a very basic workflow that uses the HDDM package for inference.

### Data

HDDM expects a dataset, provided as a `pandas DataFrame` (McKinney, 2010) with three basic columns. A `‘subj_idx’` column which identifies the subject, a `‘response’` column which specifies the choice taken in a given trial (usually coded as 1 for *correct* choices and 0 for *incorrect* choices) and a `‘rt’` column which stores the trial-wise reaction times (in seconds). Other columns can be added, for example to be used as covariates (task condition or additional measurements such as trial-wise neural data). Here we take the example of a dataset which is provided with the HDDM package. Codeblock 1 shows how to load this dataset into a Python interpreter, which looks as follows,

**Codeblock 1.**
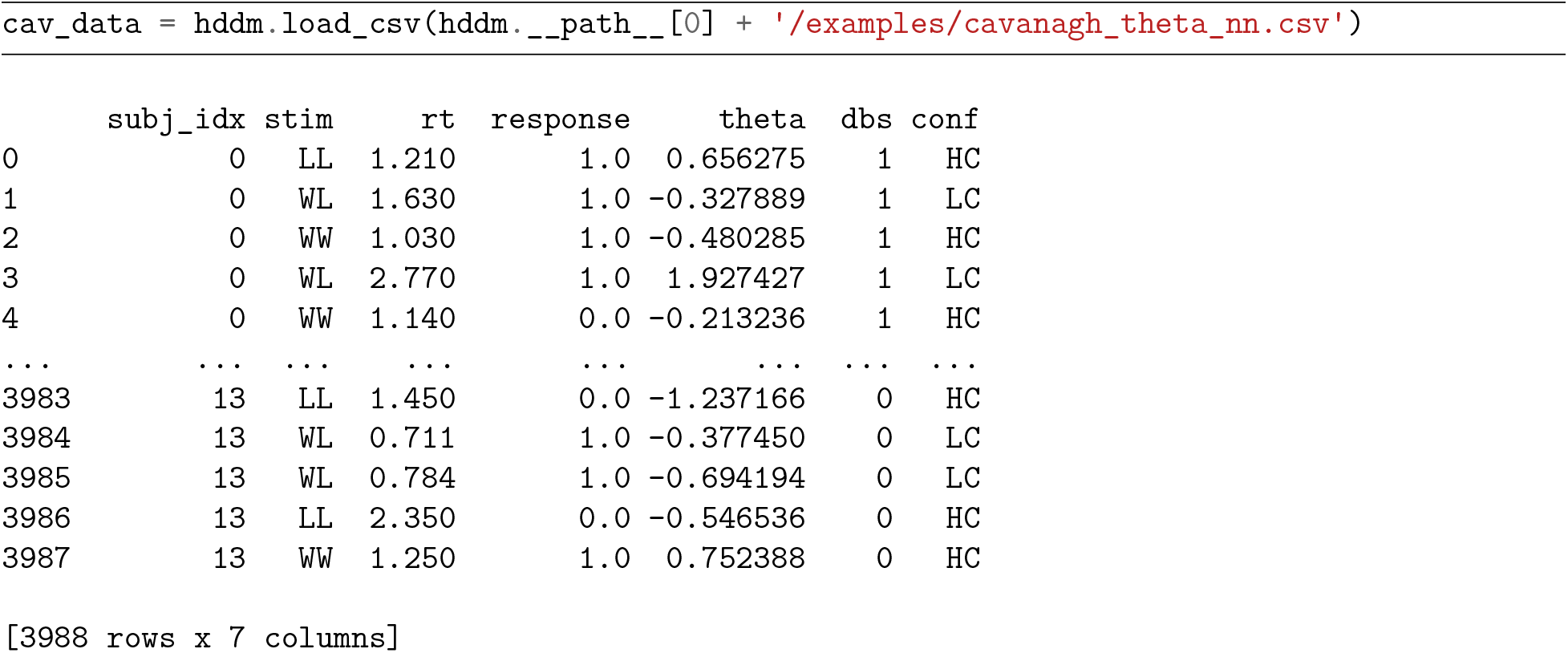
Loading package-included data

### HDDM Model

Once we have our data in the format expected by HDDM, we can now specify a HDDM model. We focus on a simple example here: a basic hierarchical model, which estimates separate drift rates (*v*) as a function of task condition, denoted by the `‘stim’` column, and moreover estimates the starting point bias *z*. (Boundary separation, otherwise known as decision threshold, *a* and non-decision time *t*, are also estimated by default).

**Codeblock 2.**
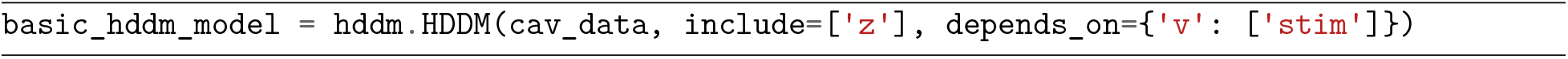
Initializing HDDM model

This model assumes that the subject level *z*, *a* and *t* parameters are each drawn from the respective group distributions, the parameters of which are also inferred. The *v* parameters derive from separate group distributions for each value of `‘stim’`. Details about the choices of *group priors* and *hyperparameters* can be found in the original toolbox paper (Wiecki et al., 2013). Codeblocks 2 and 3 show how to construct and sample from such a model.

### Sample and Analyze

With the HDDM model defined, the goal is to fit this model to a given dataset. In a Bayesian context this implies obtaining a posterior distribution over model parameters. For completeness, we note that such posterior distributions are defined via Bayes’ rule,

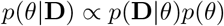

where **D** is our *data, θ* is our set of parameters, *p*(**D**|*θ*) defines the *likelihood* (analytic in the case of the standard `HDDM` class) of our dataset under the model and *p*(*θ*) defines our initial *prior* over the parameters.

HDDM uses the probabilistic programming toolbox PyMC (Patil et al., 2010) to generate samples from the posterior distribution via *Markov chain Monte Carlo* (MCMC) (specifically, using coordinate-wise slice samples (Neal, 1995)). To generate samples from the posterior we simply type,

**Codeblock 3.**
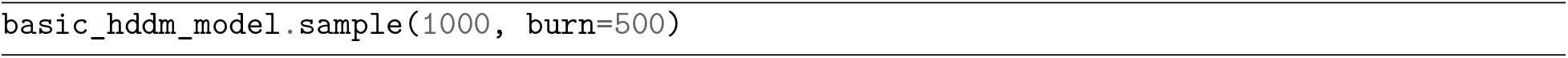
Sampling from a basic HDDM model

HDDM then provides access to a variety of tools to analyze the posterior and generate quantities of interest, including:

1. *chain summaries*: To get a quick glance at mean posterior estimates (and their uncertainty) for parameters.
2. *trace-plots* and the *Gelman-Rubin statistic* (Brooks & Gelman, 1998): To understand issues with chain-convergence (i.e., whether one can trust that the estimates are truly drawn from the posterior).
3. *the deviance information criterion* (DIC) (Spiegelhalter et al., 2014) : As a score to be used for purposes of model comparison (with caution).
4. *posterior predictive plots*: To check for the absolute fit of a given model to data (potentially as a function of task condition, etc).

The HDDM LAN extension maintains this basic HDDM workflow, which we hope facilitates seamless transition for current users of HDDM. After some brief explanations concerning *approximate likelihoods*, which form the spine of the extension, we will expose the added capabilities in detail.

## 3 Approximate Likelihoods

Approximate Bayesian inference is an active area of research. Indeed, the last decade has seen a multitude of proposals for new algorithms, many of which rely in one way or another on popular deep learning techniques (Greenberg et al., 2019; Gutmann et al., 2018; Lueckmann et al., 2019; Papamakarios & Murray, 2016; Papamakarios, Nalisnick, et al., 2019; Papamakarios, Sterratt, et al., 2019; Tejero-Cantero et al., 2020). Relevant to our goals here are algorithms which can estimate trial-by-trial likelihoods for a given model. The main idea is to replace the *likelihood* term in Bayes’ Rule, with an approximation 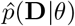, which can be evaluated via a forward pass through a simple neural network. Once the networks are trained, these “amortized” likelihoods can then be used as a plug-in (replacing the analytical likelihood function) to run approximate inference. Having access to approximate likelihoods, the user will now be able to apply HDDM to a broad variety of sequential sampling models.

The HDDM extension described here is based on a specific likelihood amortization algorithm, which we dubbed *likelihood approximation networks* (LANs) (Fengler et al., 2021). Details regarding this LAN approach, including methods, parameter recovery studies and thorough tests, can be found in Fengler et al., 2021. Note that in principle, our extension supports the integration of *any* approximate (or exact) likelihood, in the context of a now simple interface for adding models to HDDM. The scope remains limited only insofar as HDDM remains specialized towards choice / reaction time modeling. Figure 2 provides some visual intuition regarding concerning the ideas behind LANs.

**Figure 2.**
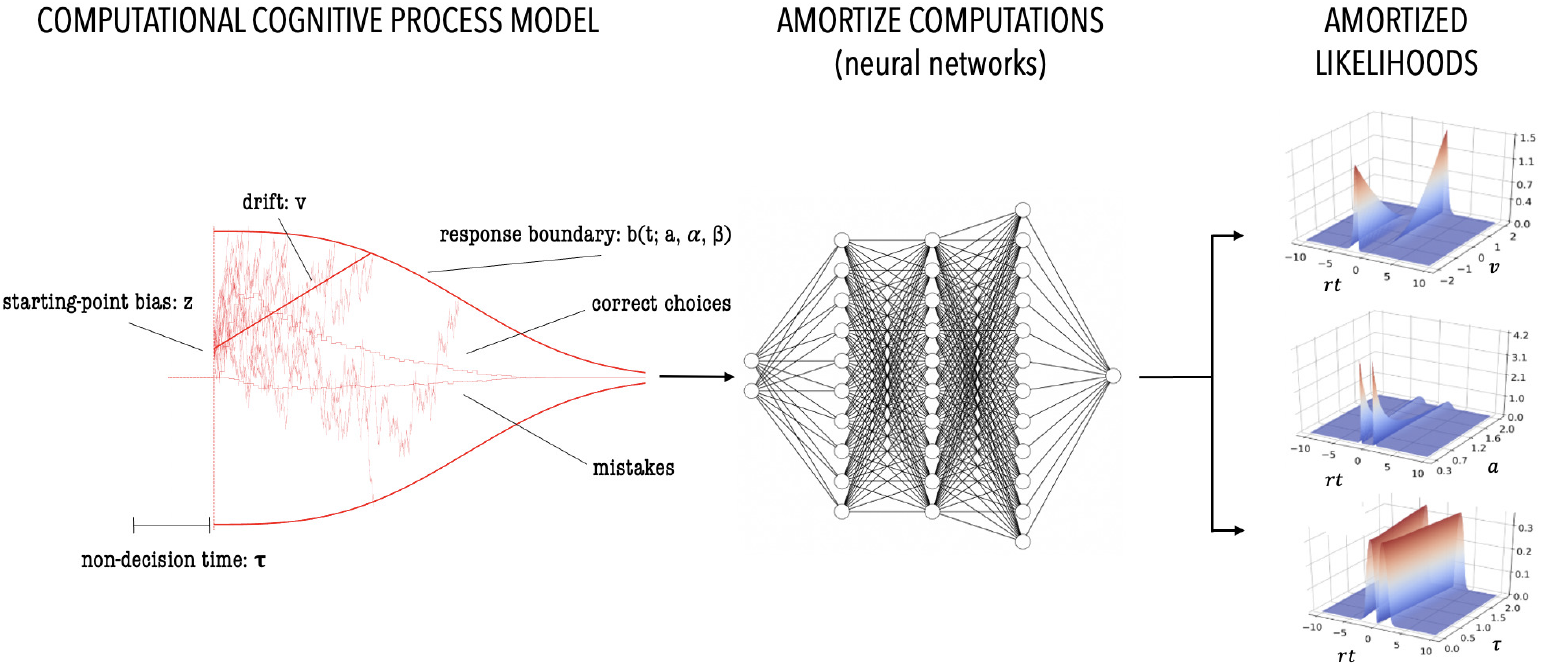
Depiction of the general idea behind *likelihood approximation networks*. We use a *simulator* of a *likelihood-free* cognitive process model to generate training data. This training data is then used to train a *neural network* which predicts the *log-likelihood* for a given feature vector consisting of model parameters, as well as a particular choice and reaction time. This neural network then acts as a stand-in for a *likelihood function* facilitating *approximate Bayesian inference*. Crucially these networks are then fully and flexibly reusable for inference on data derived from any experimental design.

## 4 HDDM Extension: Step by Step

### A Central Database For Models: hddm.model_config

To accommodate the multitude of new models, HDDM > 0.9 now uses a model specification dictionary to extract data about a given model that is relevant for inference. The `model_config` module contains a central dictionary with which the user can interrogate to inspect models that are currently supplied with HDDM. Codeblock 4 shows how to list the models included by name. For each model, we have a specification dictionary. Codeblock 5 provides an example for the simple DDM.

**Codeblock 4.**
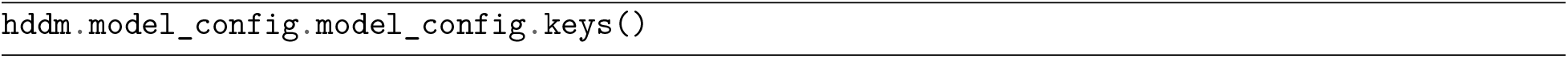
`model_config` - list available models.

**Codeblock 5.**
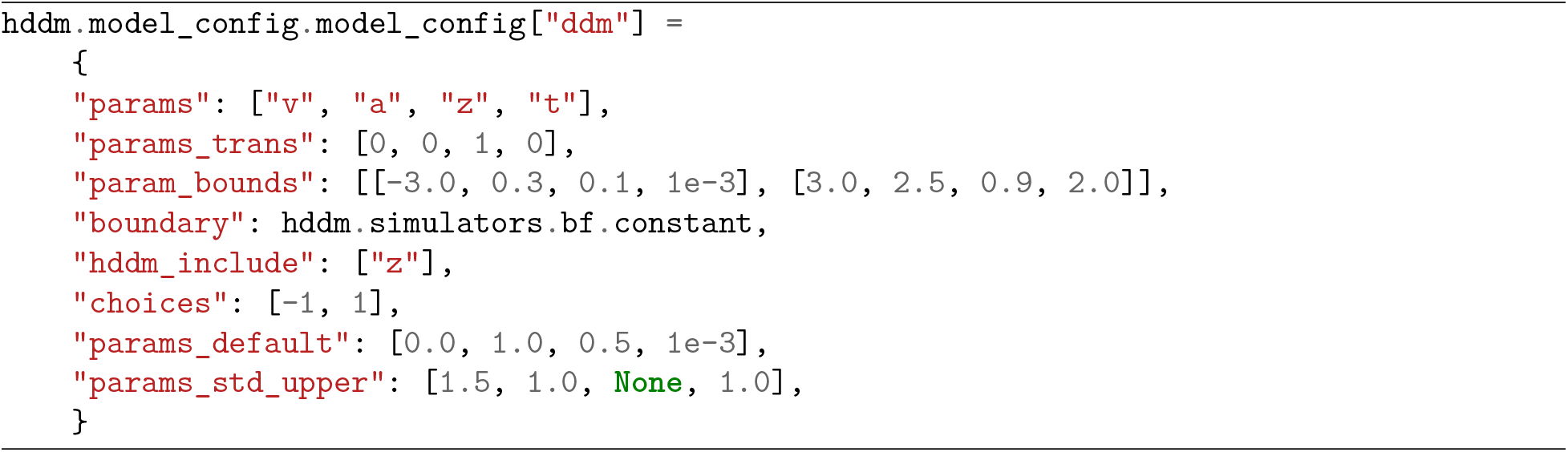
DDM specfications in `model_config`

We focus on the most important aspects of this dictionary (more options are available). Under `“params”` the parameter names for the given model are listed. `“params_trans”` specifies if the sampler should *transform* the parameter at the given position (Transforming parameters can be helpful for convergence, especially if the parameter space is strongly constrained a priori, e.g. between 0 and 1). The order follows the list supplied under `“param”`. `“param_bounds”` lists the parameter-wise lower and upper bounds of parameters that the sampler can explore. This is important in the context of LAN based likelihoods, which are only valid in the range of parameters which were observed during training. We trained the LANs included in HDDM on a broad range of parameters (spanning quite a large range of sensible data, you can inspect the training bounds in the `hddm.model_config.model_config` dictionary under the `param_bounds` key). However, it cannot be guaranteed that these were broad enough for any given empirical dataset. If the provided LANs are deemed inappropriate for a given dataset (e.g., if parameter estimates hit the bounds upon fitting), it is always possible to retrain on an even broader range of parameters. Ruling out convergence issues however, should be the first order of business in such cases.

HDDM uses the *inverse logistic* (or *logit*) transformation for the sampler to operate on an unconstrained parameter space. For a parameter *θ* and parameter bounds [*a, b*], this transformation takes *θ* from a value in [*a, b*] to a value *x* in (−∞, ∞) via,

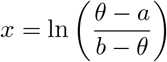

A given SSM usually has a `“decision boundary”` which is supplied as a function that can be evaluated over time-points (*t*_0_, … *t_n_*), given boundary parameters (supplied implicitly via `“params”`). The values representing each choice are reported as a `list` under `“choices”`. A note of caution: if a user wants to estimate a new model that is not currently in HDDM, a new LAN (or generally likelihood) has to be created, for it to be added to the `model_config` dictionary. Simply changing a setting in an existing `model_config` dictionary will not work. Under the `“hddm_include”` key, a list holds a working default for the `include` argument expected from the `HDDM` classes. Lastly, `“params_default”` specify the parameter values that are fixed (*not fit*) by HDDM and `“params_std_upper”` specify upper bounds on group level standard deviations for each parameter (optional, but this can help constrain the parameter ranges proposed by the sampler, making it more efficient).

These `model_config` dictionaries provide a scaffolding for model specification which is applied throughout all of the new functionalities discussed in the next sections.

### Batteries Included: hddm.simulators, hddm.network_inspectors

The new `HDDMnn` (where *nn* is for *neural network*), `HDDMnnRegressor` and `HDDMnnStimCoding` classes have access to a (growing) stock of supplied SSMs, including rapid compiled (Behnel et al., 2010) simulators, and rapid likelihood evaluation via LANs (Fengler et al., 2021) and their implementation in PyTorch (Paszke et al., 2019). We will discuss how to fit these models to data in the next section. Here we describe how one can access the low level simulators and LANs directly, in case one wants to adopt them for custom purposes. We also show how to assess the degree to which the LAN approximates the true (empirical) likelihood for a given model. Users who just want to apply existing SSMs in HDDM to fit data can skip to the next section.

As described in the previous section, the user can check which models are currently available by using the `model_config` dictionary. Figure 3, provides some pictorial examples. For a given model, a doc-string includes some information (and possible warnings) about usage. As an example, let us pick the **angle** model, which is a SSM that allows for the decision boundary to decline linearly across time with some estimated angle (note that although other aspects of the model are standard DDM, even in this case the likelihood is analytically intractable. Nevertheless, we previously observed that inference using LANs yields good parameter recovery, as per Fengler et al., 2021).

**Figure 3.**
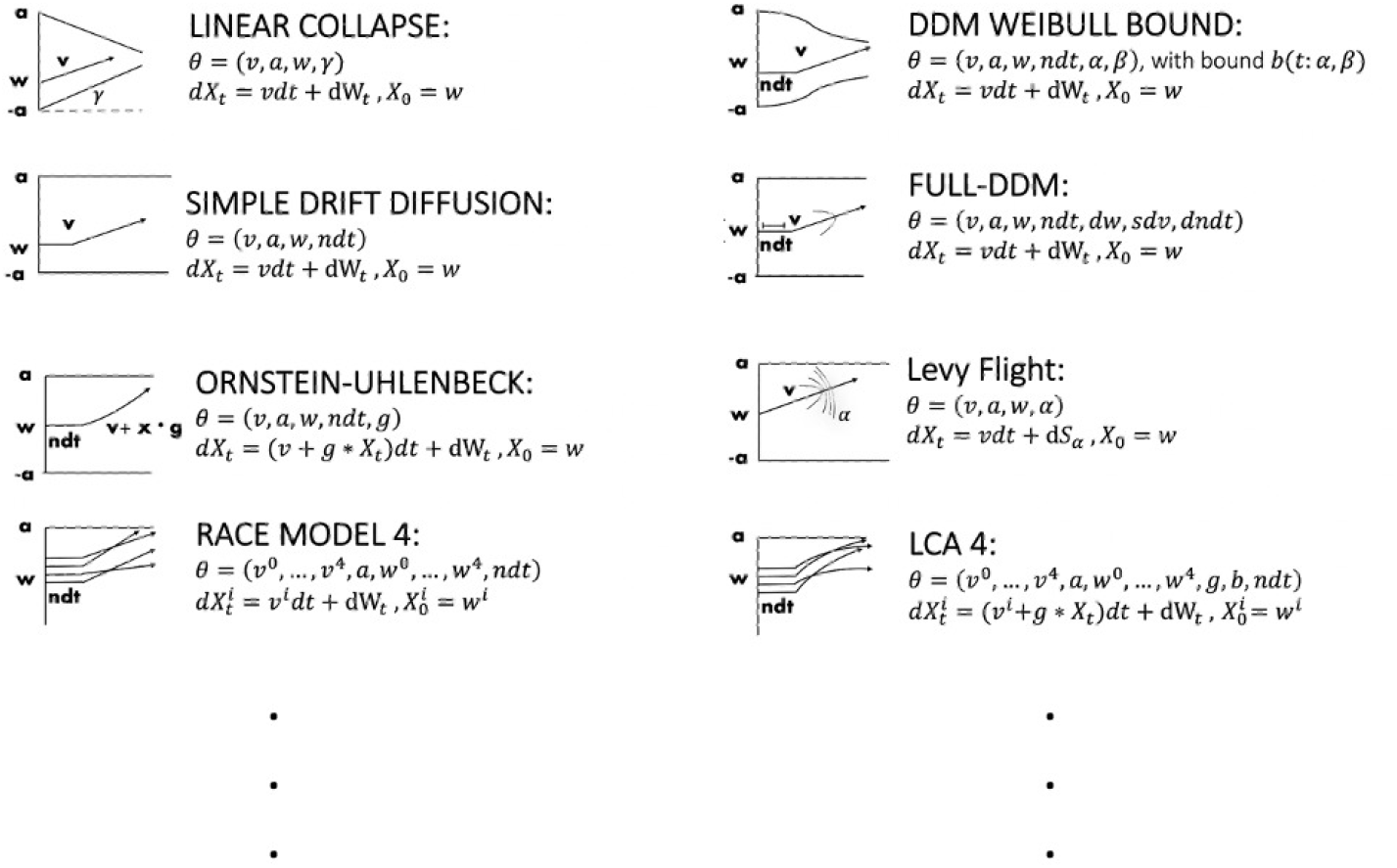
Graphical examples for some of the sequential sampling included in HDDM.

Codeblock 6, illustrates such a doc-string. Codeblock 7 shows how we can simulate synthetic data from this model. Following the code, the variable out is now a three-tuple. The first element contains an array of *reaction times*, the second contains an array of *choices* and finally the third element returns a *dictionary of metadata* concerning the simulation run. Next, we can access the LAN corresponding to our **angle** model directly by typing the code in codeblock 8. We can utilize the `get_torch_mlp` function, which is defined in the `network_inspectors` submodule.

**Codeblock 6.**
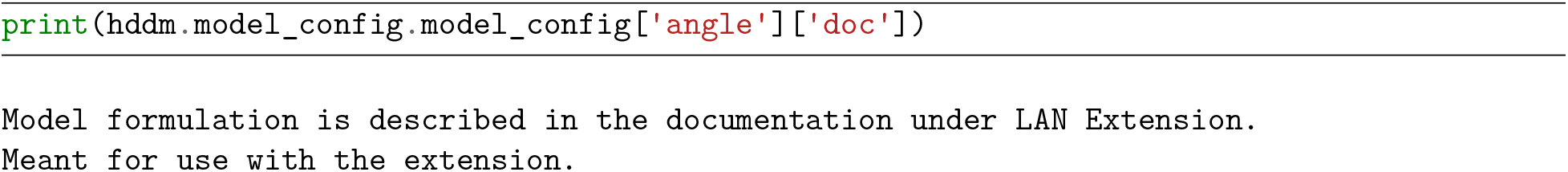
`model_config` doc-string for the `angle` model

**Codeblock 7.**
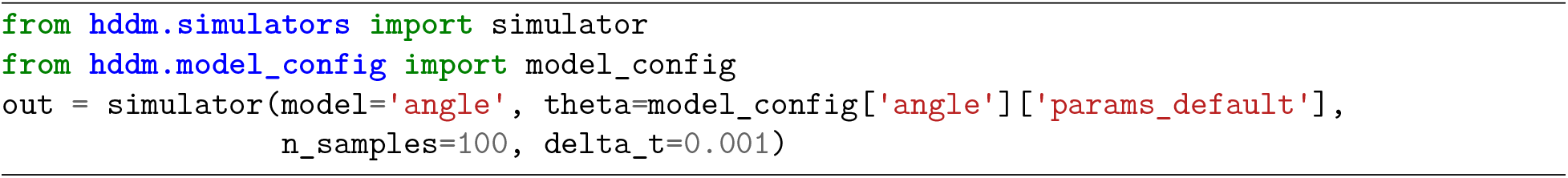
Using the `simulator` simulator for generating synthetic data

**Codeblock 8.**
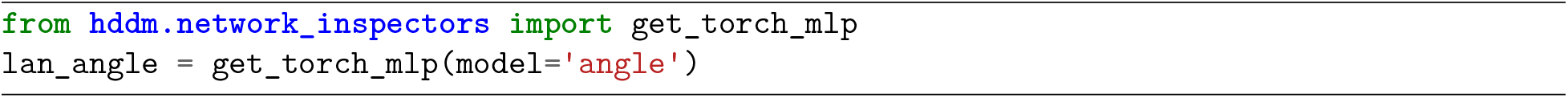
Loading a torch network from the package

The `lan_angle` object defined in codeblock 8 is in fact a method, which defines the forward pass through a given LAN. It expects as input a matrix where each row defines a parameter vector suitable for the SSM of choice (here **angle**, so we need a value for each of the parameters `[‘v’, ‘a’, ‘z’, ‘t’, ‘theta’]` which can be found in our `model_config` dictionary). Two elements are then added: a *reaction time* and a *choice* at which we would like to evaluate our likelihood. Codeblock 9 provides a full example.

To facilitate a simple sanity check, we provide the `kde_vs_lan_likelihoods` plot, which can be accessed from the `network_inspectors` submodule. This plot lets the user compare LAN likelihoods against empirical likelihoods from simulator data for a given matrix of parameter vectors (Fengler et al., 2021). The empirical likelihoods are defined via kernel density estimators (KDEs) (Silverman, 1986). We show an example in codeblock 10. Figure 4 shows the output.

**Figure 4.**
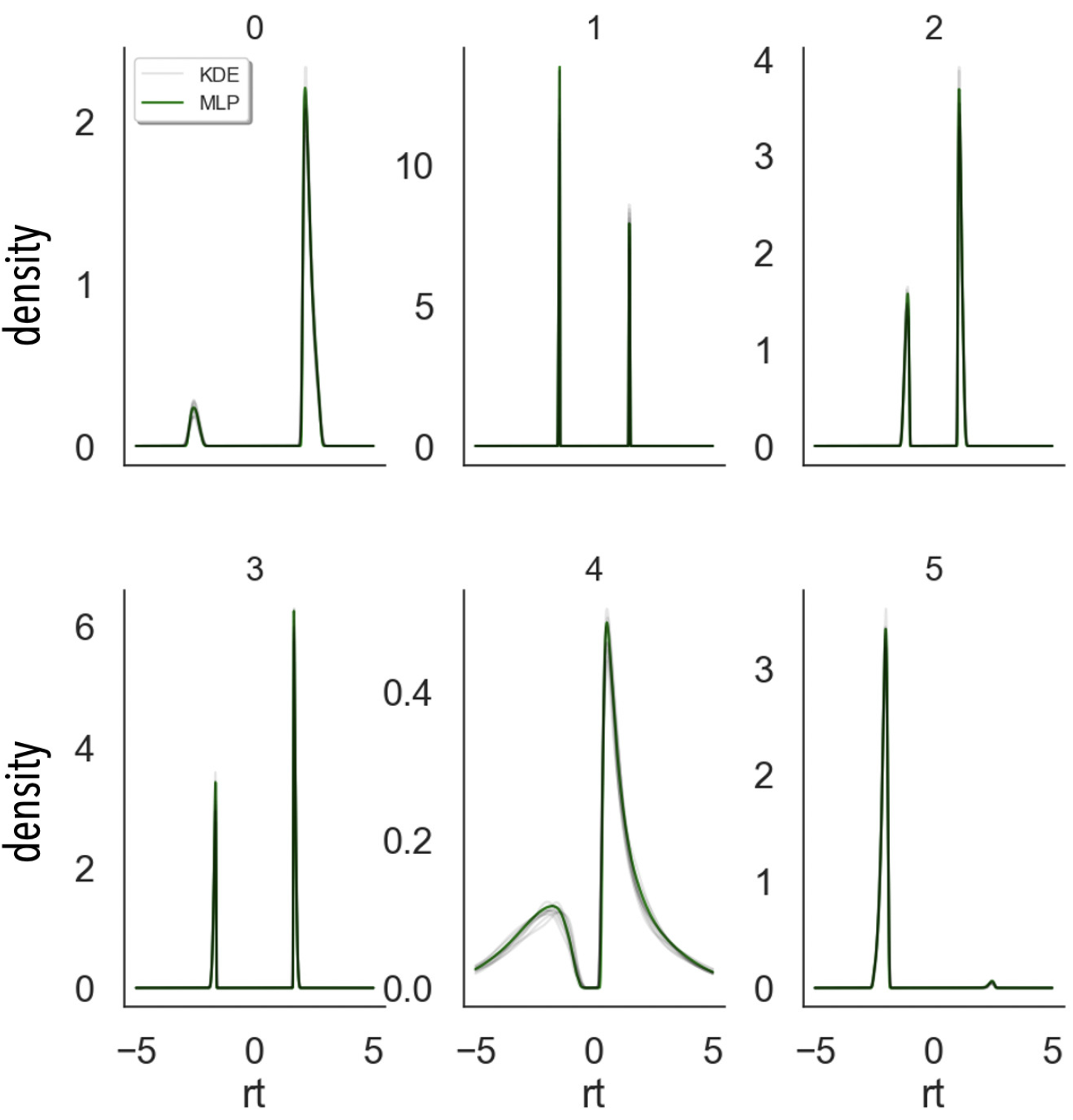
Example of a `kde_vs_lan_likelihoods` plot. If the green (deterministic) and gray (stochastic) lines overlap, then the approximate likelihood (MLP for multilayered perceptron, the neural network that provides our LAN) is a good fit to the actual likelihood.

**Codeblock 9.**
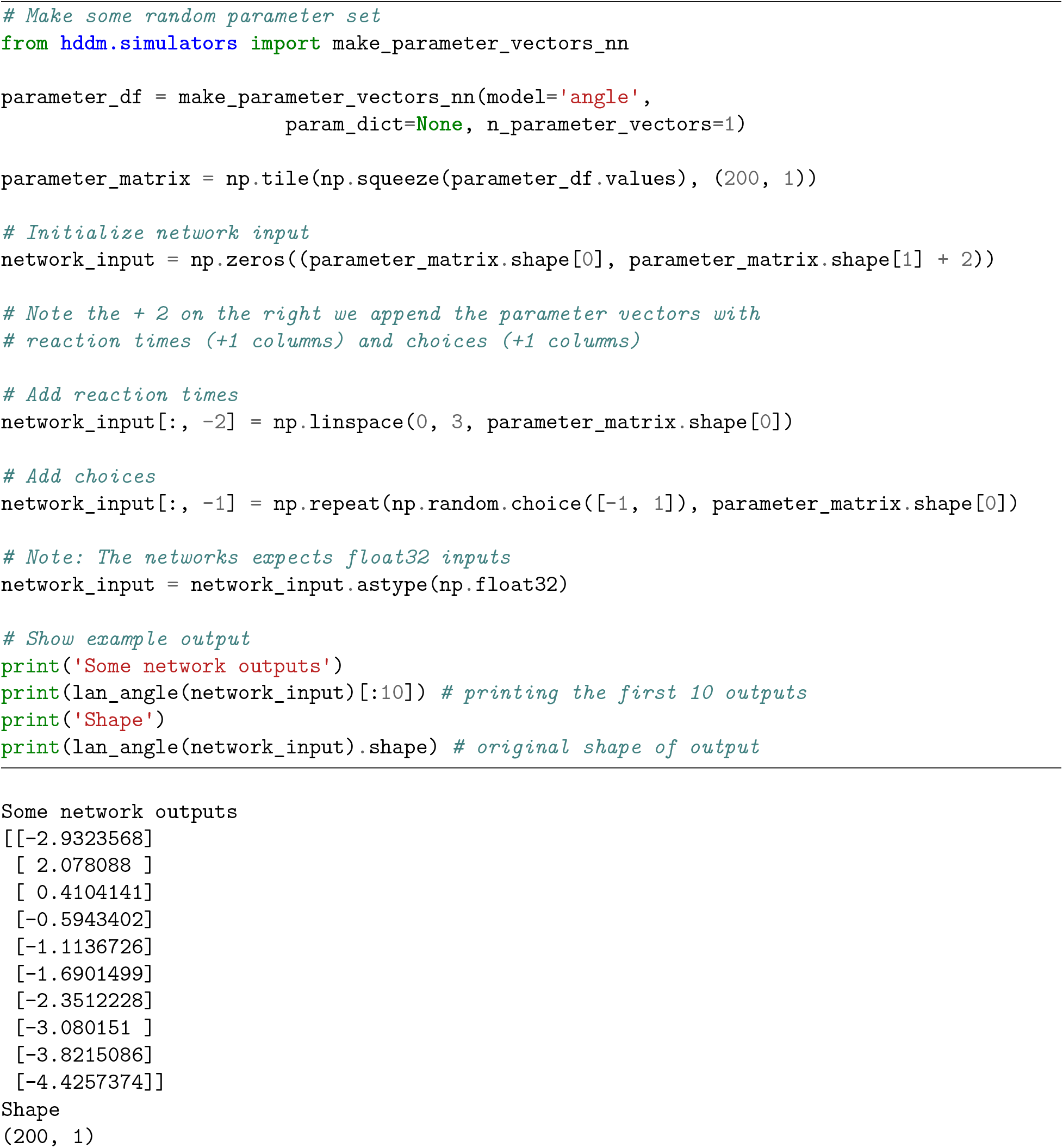
Check forward pass of supplied `angle` network.

### Fitting data using HDDMnn, HDDMnnRegressor, and HDDMnnStimCoding classes

Using the `HDDMnn`, `HDDMnnRegressor` and `HDDMnnStimCoding` classes, we can follow the general workflow established by the basic `HDDM` package to perform Bayesian inference. In this section we will fit the **angle** model to the example dataset provided with the HDDM package. Codeblock 11 shows us how to load the correponding dataset, after which we can set up our HDDM model, and draw 1000 MCMC samples using the code in codeblock 12.

**Codeblock 10.**
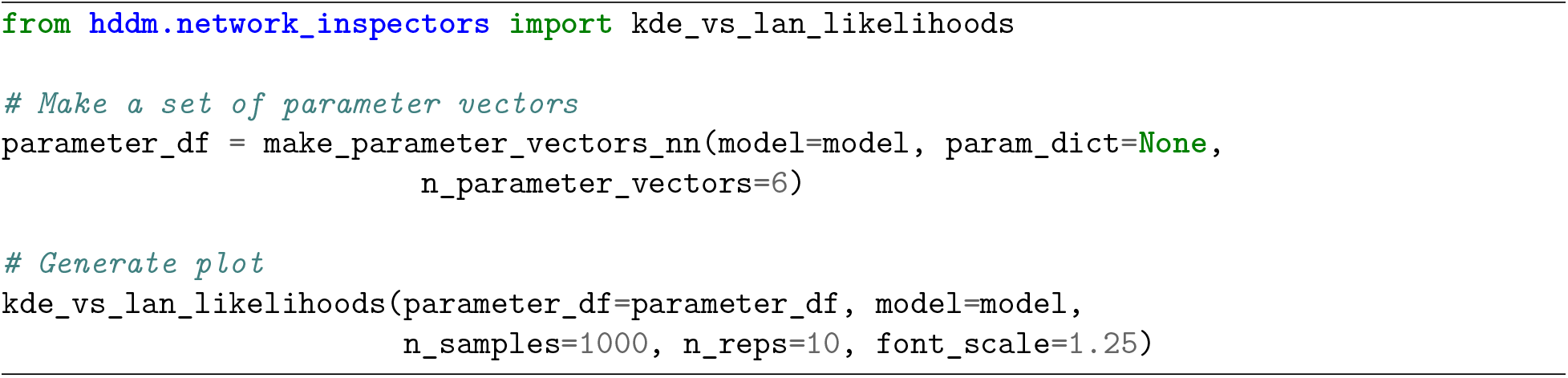
Example usage of the `kde_vs_lan_likelihood()` function to compare LAN likelihoods to empirical kernel-density estimates.

We note a few differences between a call to construct a `HDDMnn` class and a standard `HDDM` class. First, the supply of the `model` argument specifying which SSM to fit (requires that this model is already available in HDDM; see above). Second, the inclusion of model-specific parameters under the `include` argument. The workflow is otherwise equivalent, a fact that is conserved for the `HDDMnnRegressor` and `HDDMnnStimCoding` classes. A third difference concerns the choice of argument defaults. The `HDDMnn` class uses non-informative priors, instead of the informative priors derived from the literature which form the default for the basic `HDDM` class. Since, as per our earlier discussions, variants of SSMs are historically rarely fit to experimental data, we can not easily derive reasonable informative priors from the literature and therefore choose to remain agnostic in our beliefs about the parameters underlying a given dataset. If the research community starts fitting SSM variants to experimental data, this state of affairs may evolve through collective learning. At this point we caution the user to however not use these new models blindly. We strongly encourage conducting appropriate parameter recovery studies, specific to the experimental dataset under consideration. We refer to the section on *inference validation tools* below, for how HDDM might help in this procedure.

**Codeblock 11.**
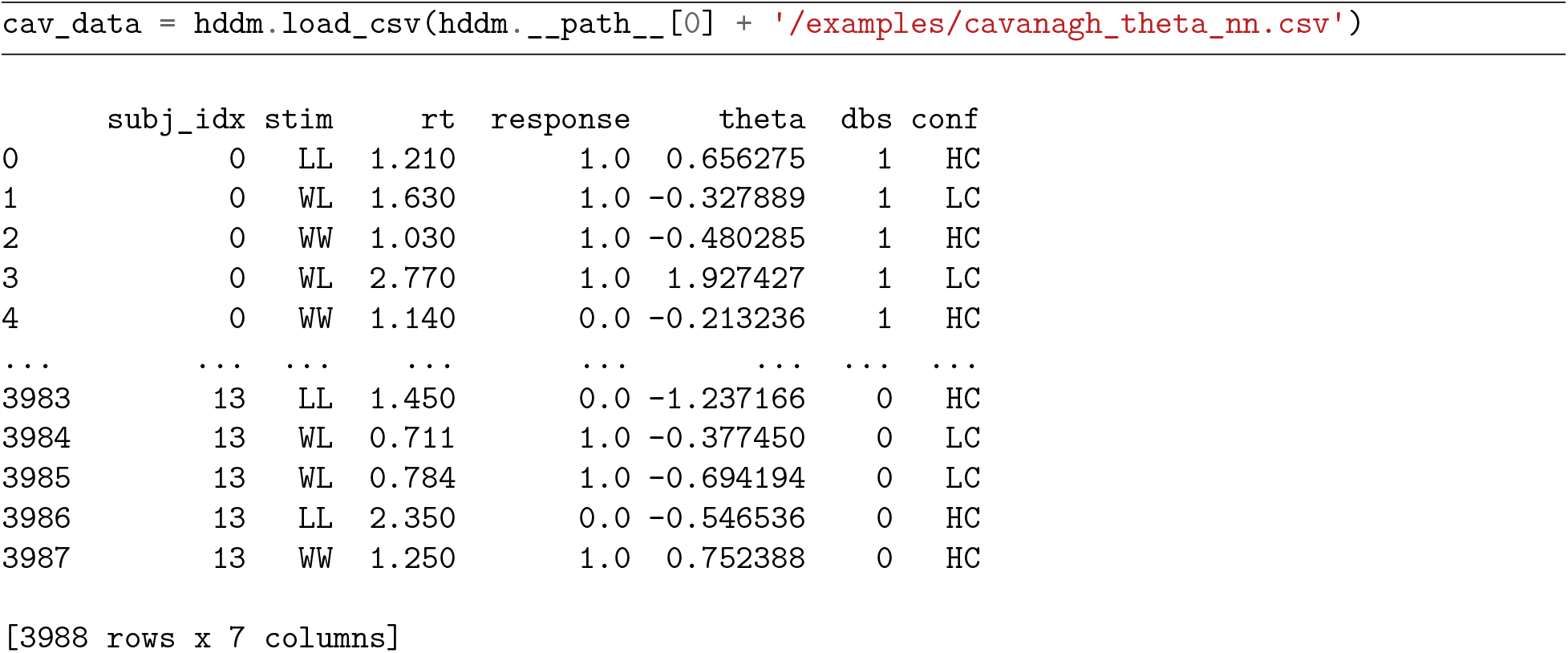
Loading package supplied **cavanagh** dataset.

**Codeblock 12.**
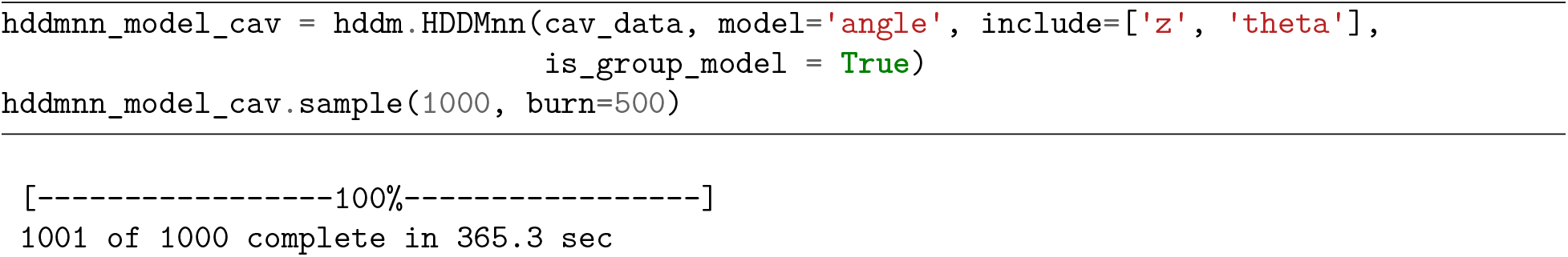
Sampling from a HDDMnn model.

### New Visualization Plots: hddm.plotting

Based on our model fit from the previous section, we illustrate a few new informative plots, which are now included in HDDM. We can generally distinguish between two types of plots. Plots which use the traces only (to display posterior parameter estimates) and plots which make use of the model simulators (to display how well the model can reproduce empirical data given posterior parameters). The first such plot is produced by the the `plot_caterpillar` function, which presents an approximate posterior 99%-HDI (specifically we show the 1% to 99% range in the cumulative distribution function of the posterior), for each parameter. Codeblock 13 shows us how to invoke this function and Figure 5 illustrates the resulting plot.

**Figure 5.**
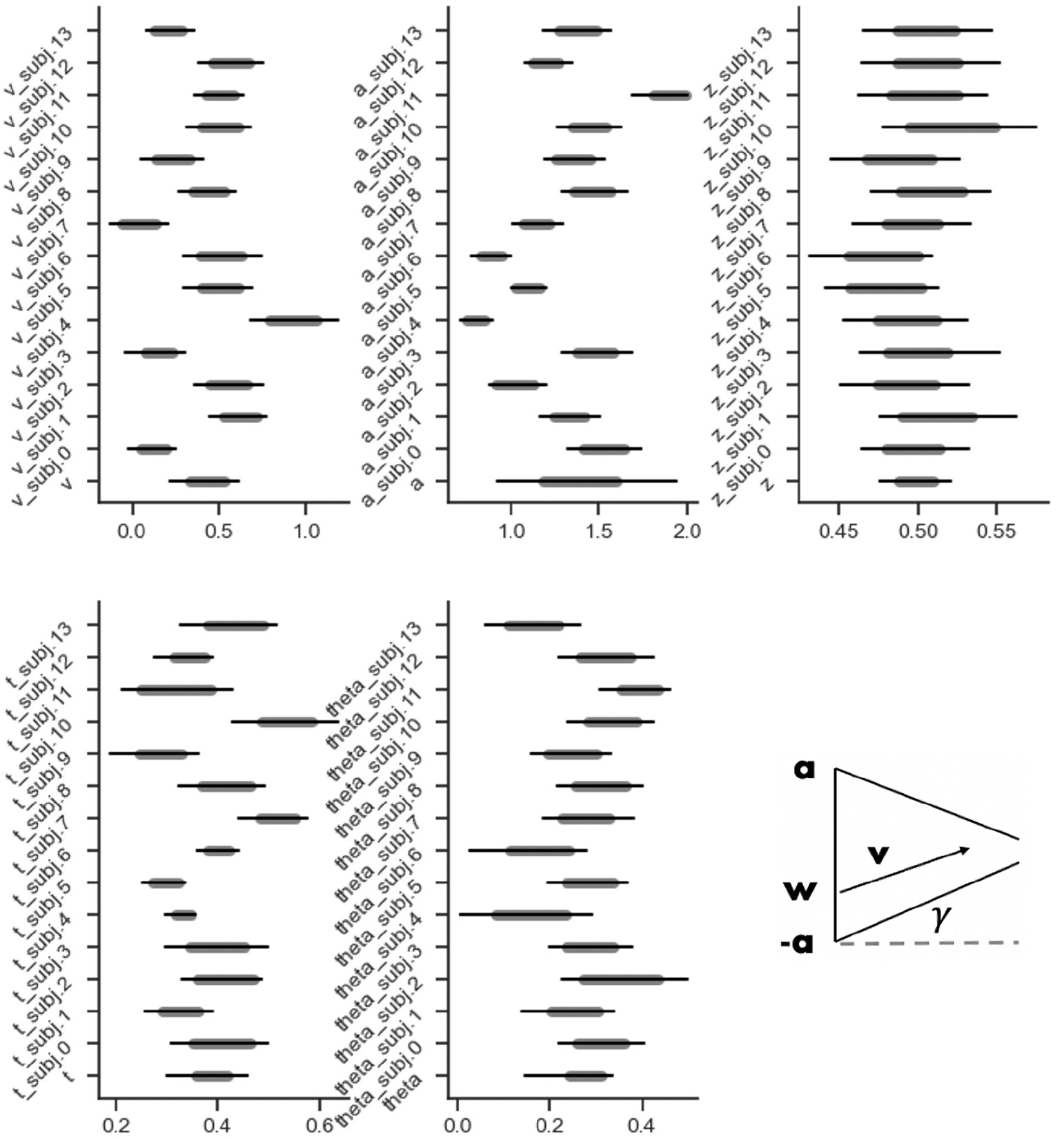
Example of a `caterpillar_plot`. The plot, split by model parameters, shows the 99% (line-ends) and 95% (gray band ends) highest density intervals (HDIs) of the posterior for each parameter. Multiple styling options exist.

The second such plot is the a posterior pair plot, called via the `plot_posterior_pair` function. This plot shows the pairwise posterior distribution, subject by subject (and, if provided, condition by condition). Codeblock 14 illustrates how to call this function, and Figure 6 exemplifies the resulting output.

**Figure 6.**
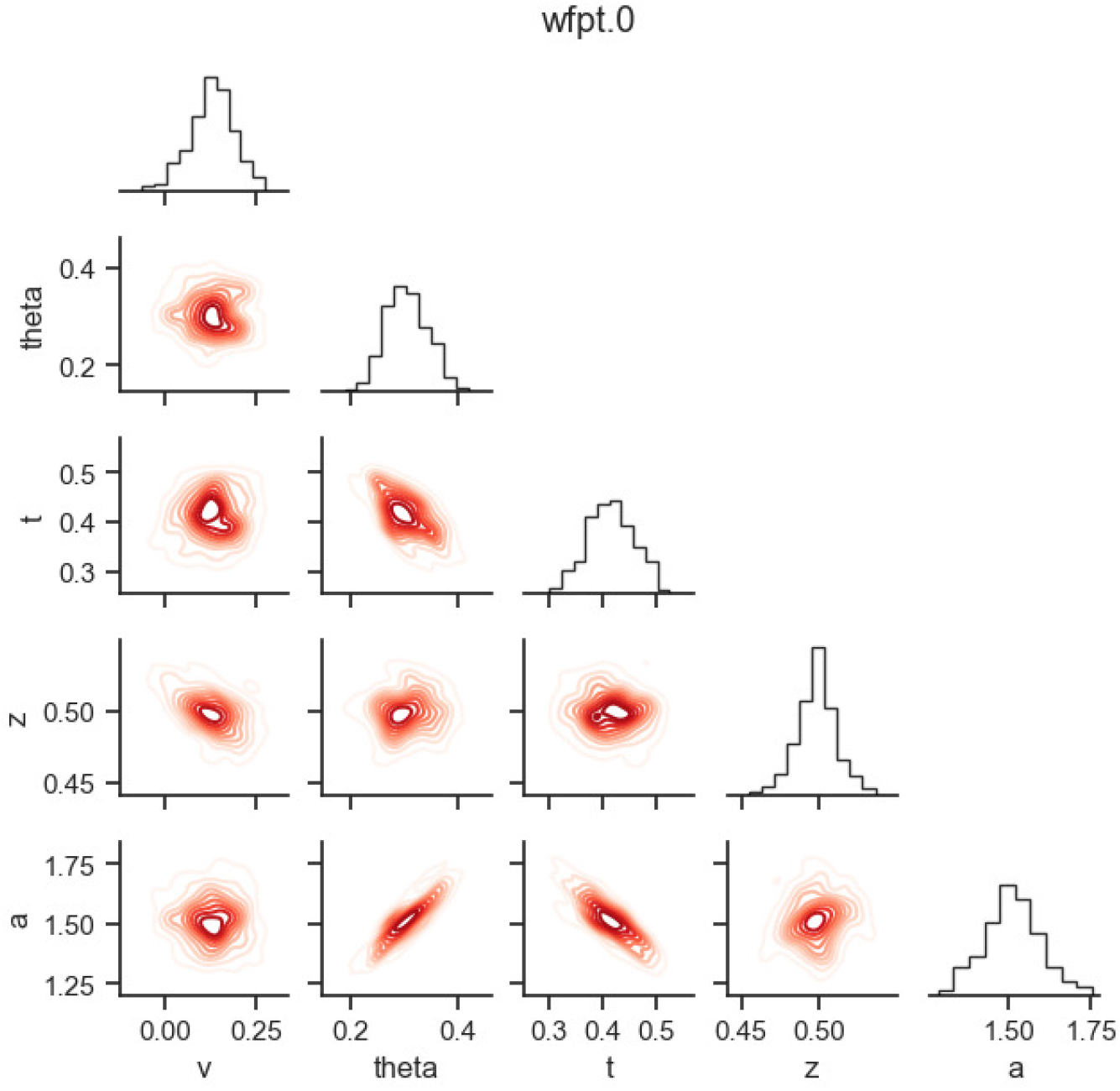
Example of a `posterior_pair_plot` in the context of parameter recovery. The plot is organized per stochastic node (here, grouped by the `‘subj_idx’` column where in this example `‘subj_idx’ = ‘0’`). The diagonal shows the *marginal posterior* of a given parameter as a histogram. The elements below the diagonal show pair-wise posteriors via (approximate) level curves. These plots are especially useful to identify parameter collinearities, which indicate parameter-tradeoffs and can hint at issues with identifiability. This example shows how the `theta` (boundary collapse) and `a` (boundary separation) parameters as well as the `t` (non-decision time) and `a` parameters trade-off in the posterior. We refer to Fengler et al., 2021 for parameter recovery results using the underlying *angle* SSM. We note that such parameter trade-offs and attached identifiability issues derive not just from a given likelihood model, but are also affected by the data and parameter structure as task design and modeling choices.

A last very useful plot addition is what we call the **model plot**, an extension to the standard posterior predictive plot, which can be used to visualize the impact of the parameter posteriors on decision dynamics. For example, if one is estimating a linearly collapsing bound, instead of just interpreting the posterior angle parameter, one can see how that translates to the evolving decision bound over time in tandem with the estimating drift rate, etc. It is an extension of the `plot_posterior_predictive` function. This function operates by manipulating `matplotlib axes` objects, via a supplied *axes manipulator*. The novel *axis manipulator* in the example show in Codeblock 15 is the `_plot_func_model` function. Figure 7 show the resulting plot.

**Figure 7.**
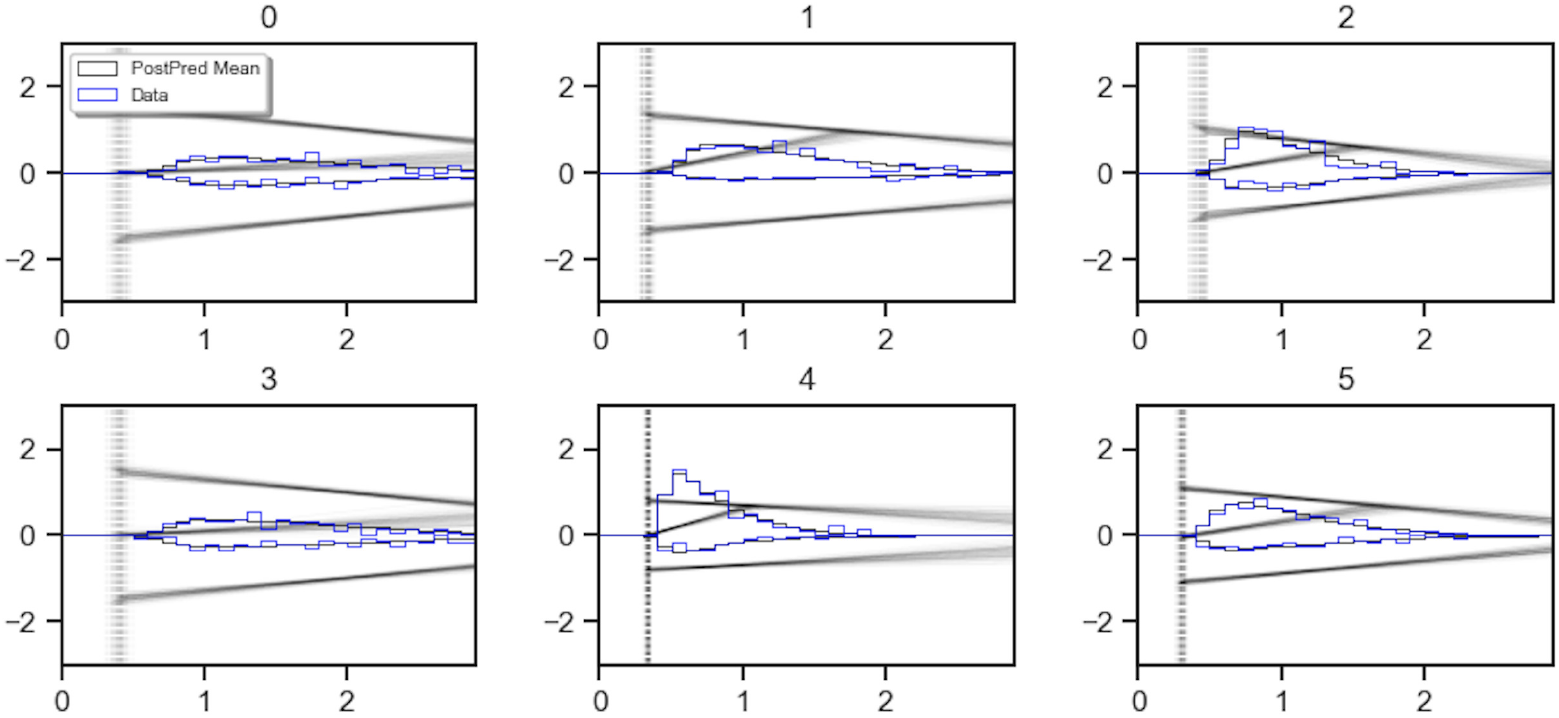
Example of a `model_plot`. This plot shows the underlying data in blue, choices and reaction times presented as histograms (positive y-axis for choice option 1, negative y-axis or choice option 0 or −1). The black histograms show the reaction times and choices under the parameters corresponding to the posterior mean. In addition the plot shows a graphical depiction of the model corresponding to parameters drawn from the posterior distribution in black. Various options exist to add and drop elements from this plot; the provided example corresponds to what we consider the most useful settings for purposes of illustration. Note that, in the interest of space, we only illustrate the first six subjects here.

**Codeblock 13.**
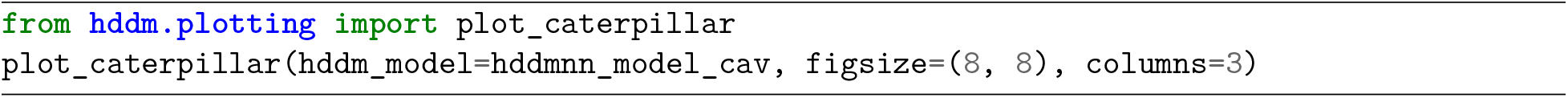
Example usage of the `caterpillar_plot()` function.

**Codeblock 14.**
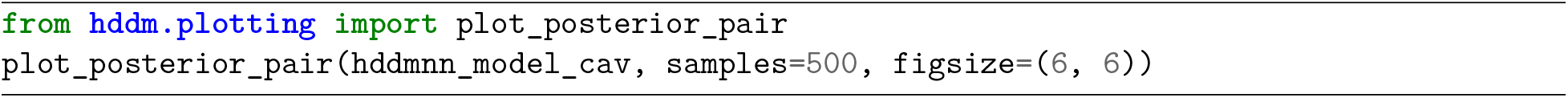
Example concerning usage of the `plot_posterior_pair()` function.

We use this moment to illustrate how the **angle** model in fact outperforms the **DDM** on this example dataset. For this purpose we take an example subject from Figure 7 and contrast the posterior predictive of the **angle** model with the posterior predictive of the **DDM** side by side in Figure 8. We clearly see that the **DDM** model has trouble capturing the leading edge and the tail behavior of the rt distributions simultaneously, while the **angle** model strikes a much better balance. While this example does not present a fully rigorous model comparison (DIC scores for example however bear out the same conclusion) exercise, it provides a hint at the benefits one may expect from utilizing an expanded model space.

**Figure 8.**
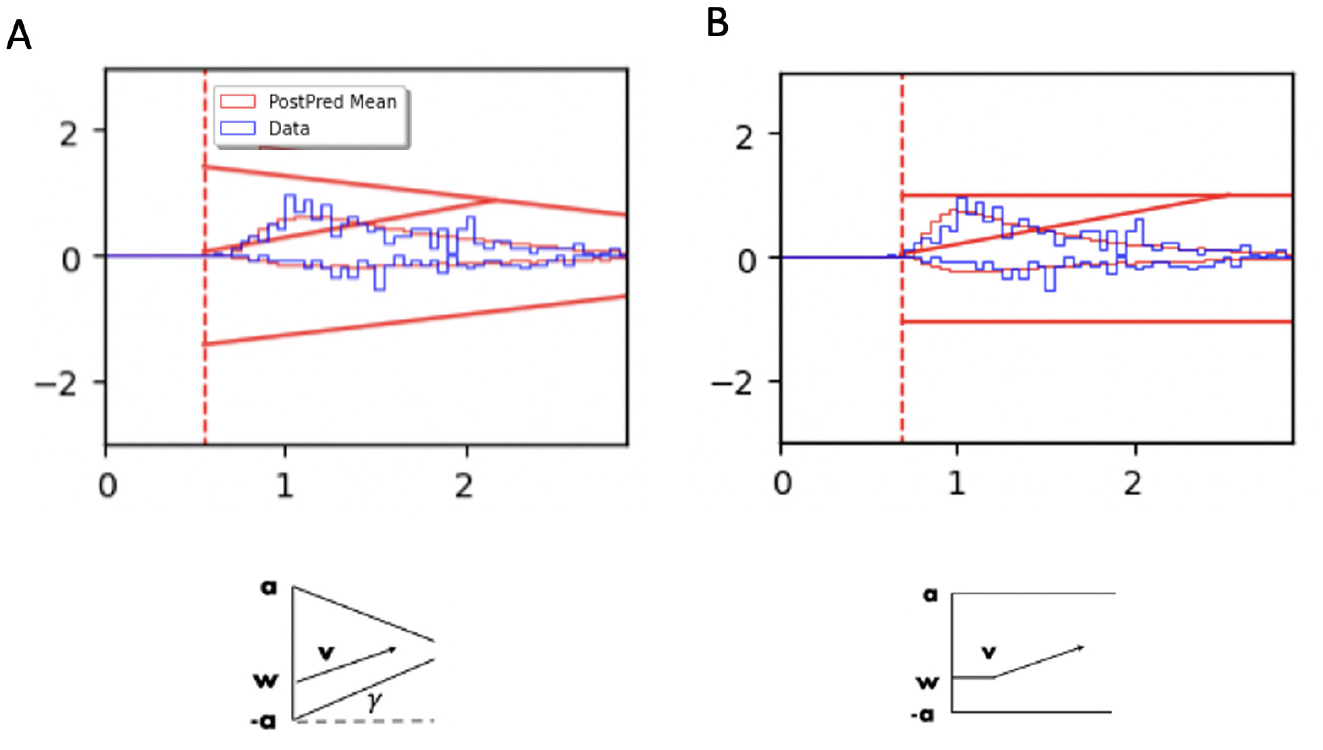
Contrasting the posterior predictive of the **angle** and **DDM** model on an example subject. **A**) shows the **angle** model and **B**) shows the **DDM**. While the fits are not dramatically better for this dataset (in our experience more extreme differences can be seen in other cases), the **angle** model shows two characteristic differences to the **DDM** model fit. First, it better captures the graceful initial increase in density for short reaction times. Second, it captures the slower decrease in density for longer reaction times, as compared to the **DDM**, for which the reaction time density falls of quicker than is apparent in the data. Both of these effects are directly produced by allowing a collapsing bound, instead of the **DDM’s** static, parallel bounds.

**Codeblock 15.**
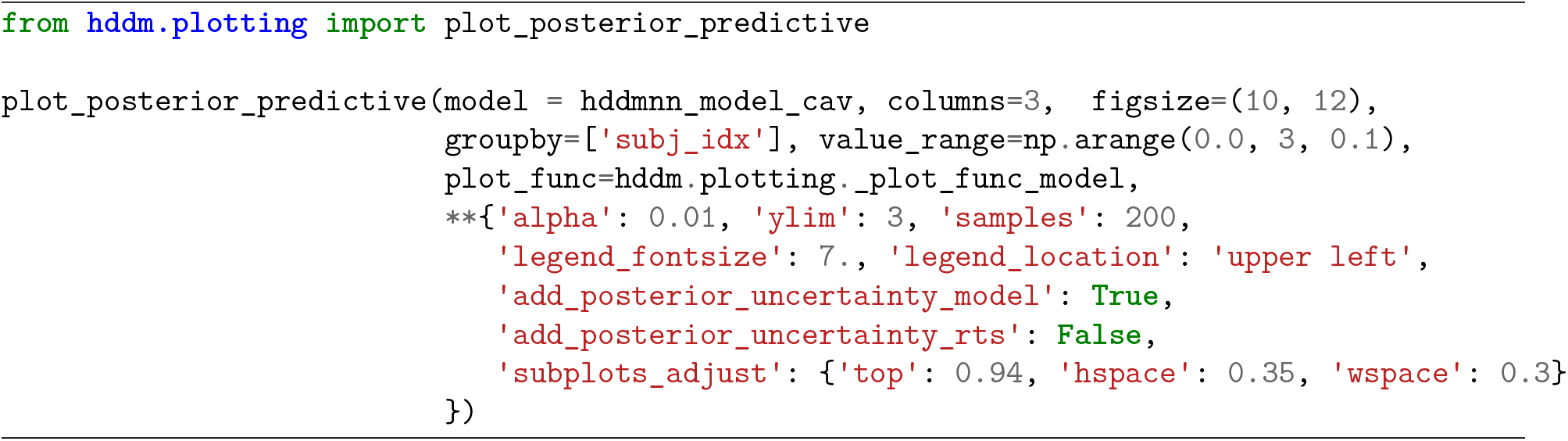
Example usage of the `plot_posterior_predictive()` function

### Inference Validation Tools: simulator_h_c()

Validating that a model is identifiable on simulated data is an important aspect of a trustworthy inference procedure (Evans et al., 2020; Holmes & Trueblood, 2018; Tran et al., 2021; Wilson & Collins, 2019). We have two layers of uncertainty in this regard. First, LANs are approximate likelihoods. A model that is otherwise identifiable could in principle lose this property when using LANs to estimate its parameters from a dataset, should the LAN not have been trained adequately. Second, a given model can inherently be unidentifiable for a given dataset and or theoretical commitments (regardless of whether its likelihood is analytic or approximate). As a simple example consider an experimental dataset, which does not include enough samples to identify the parameter of a model of interest with any degree of accuracy. Slightly more involved, the posterior could tend to be multi-modal, a problem for MCMC samplers that can lead to faulty inference. While increasing the number of trials in an experiment and/or increasing the number of participants can help remedy this situation, this is not a guarantee. Apart from the size and structure of the empirical dataset, our modeling commitments play an important role for identifiability too. As an example, we might have experimental data from a random dot motion task and we are interested in modeling the *choices* and *reaction times* of participating subjects with our **angle** model. A reasonable assumption is that the *v* parameter (a rough proxy for processing speed) differs depending on the difficulty of the trial. However, the parameters *t* and *a* may not depend on the difficulty since we do not have a good a-priori theoretical reason to suspect that the *non-decision time* (*t*) and the initial the *boundary separation a* (the degree of evidence expected to take a decision) will differ across experimental conditions. These commitments are embedded in the model itself (they are assumptions on the data generating process imposed by the modeler), and determine jointly with an experimental dataset whether inference can be successful. For a modeler it is therefore of paramount importance to check whether their chosen combinations of theoretical commitments and experimental dataset jointly lead to an inference procedure that is accurate. Since the space of models incorporated into HDDM has been significantly expanded with the LAN extension, we provide a few tools to help facilitate parameter recovery studies which are relevant to real experimental data analysis and plan to supplement these tools even further in the future.

First, we provide the `simulator_h_c` function, in the `hddm_dataset_generators` submodule. The function is quite flexible, however we will showcase a particularly relevant use-case. Taking our `cav_data` dataset loaded previously, we would like to generate data from our **angle** model in such a way that we encode assumptions about our model into the generated dataset. In the example below we assume that the *v* and *theta* parameters vary as a function of the `“stim”` column. For each value of `“stim”` a group-level *μ* and *σ* (defining the *mean* and *standard deviation* of a group level Normal distribution) are generated and subject-level parameters are sampled from this group distribution. This mirrors exactly the modeling assumptions when specifying a HDDM model with the `depends_on` argument set to `{‘v’ : ‘stim’, ‘theta’ : ‘stim’}`. Codeblock 16 provides an example on how to call this function. The `simulator_h_c` function returns the respective dataset (here `sim_data`) exchanging values in the previous *rt* and *response* columns with simulation data. Trial-by-trial parameters are attached to the dataframe as well. The `parameter_dict` dictionary contains all the parameters of the respective hierarchical model which was used to generate the synthetic data. This parameter dictionary follows the parameter naming conventions of HDDM exactly. We can fit this data using the `HDDMnn` class as illustrated in Codeblock 17.

**Codeblock 16.**
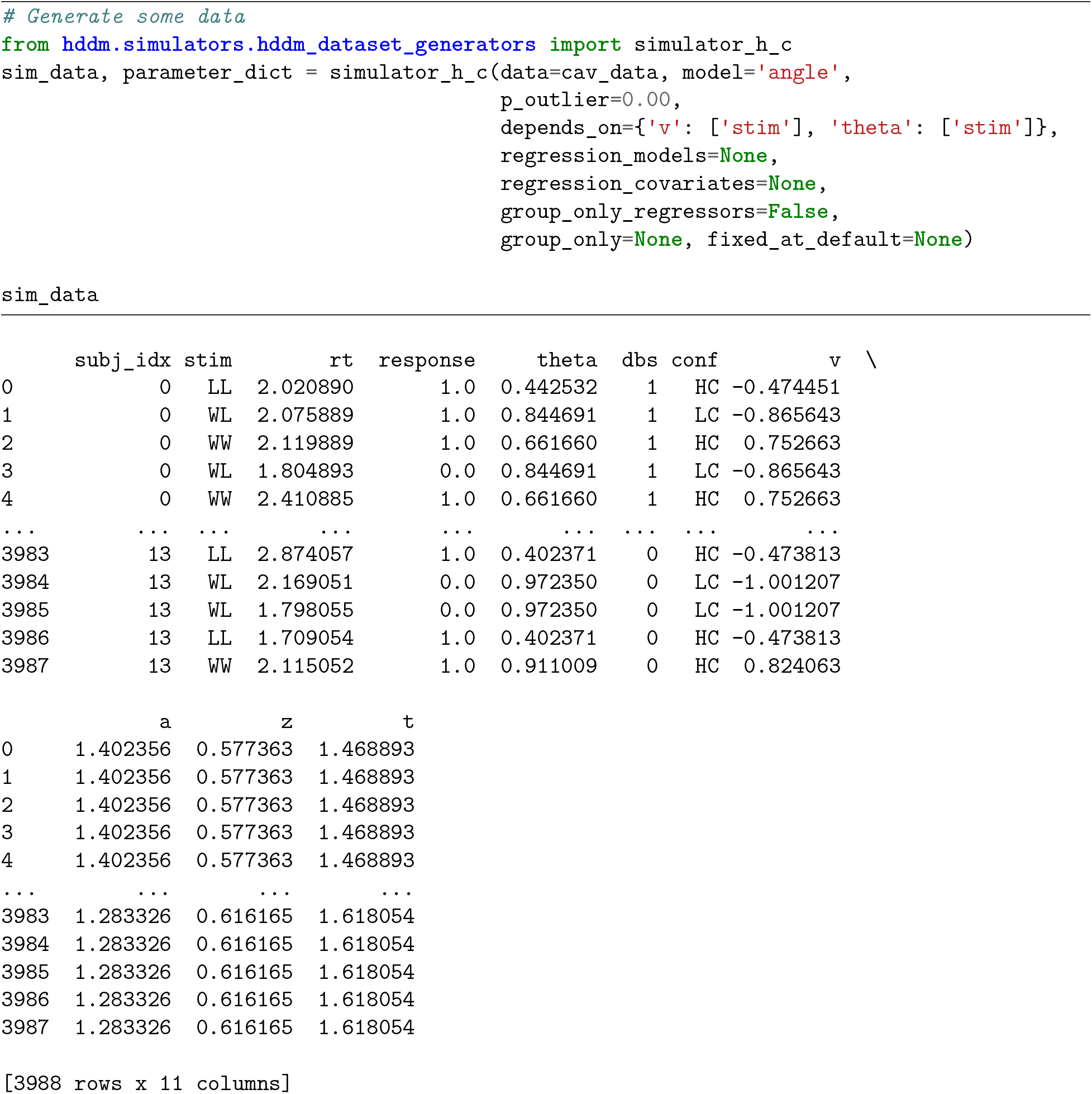
Using the `simulator_h_c()` function.

The plots defined in the previous section allow us to specify a `parameter_recovery_mode` which we can utilize to check how well our estimation worked on our synthetic dataset. Codeblocks 18, 19 and 20 and Figures 9, 10 and 11 show respectively code and plot examples.

**Figure 9.**
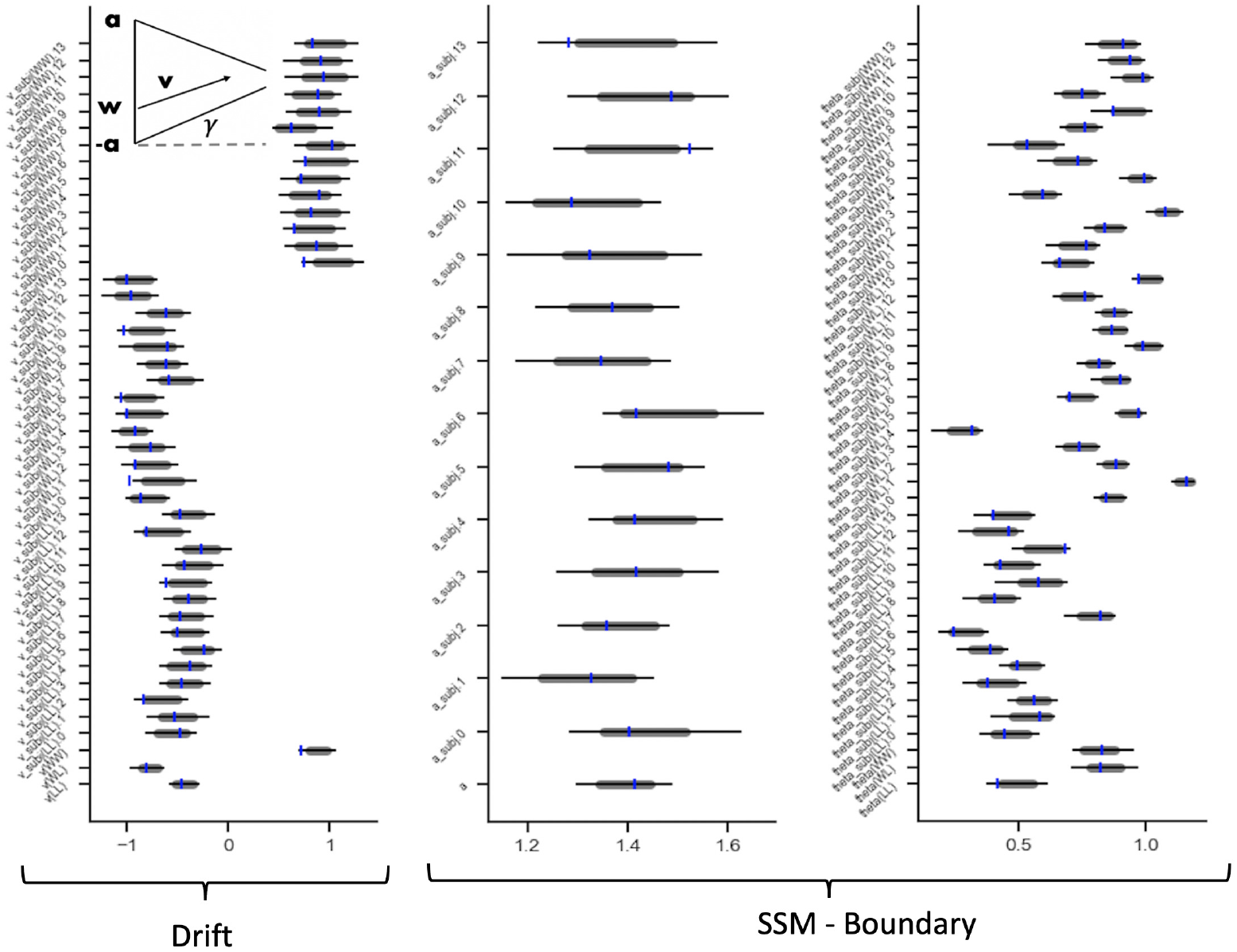
Example of a `caterpillar_plot`. The plot is split by model parameter kind, showing parameter-wise, the 99% (line-ends) and 95% (gray band ends) highest density intervals (HDIs) of the posterior. In the context of parameter recovery studies, the user can provide ground-truth parameters to the plot, which will be shown as blue tick-marks on top of the HDIs. Multiple styling options exist. Note, in the interest of space, we show only three of the five basic parameters here. `[‘v’, ‘a’, ‘theta’]` of the underlying model (leaving out `[‘z’, ‘t’]`).

**Figure 10.**
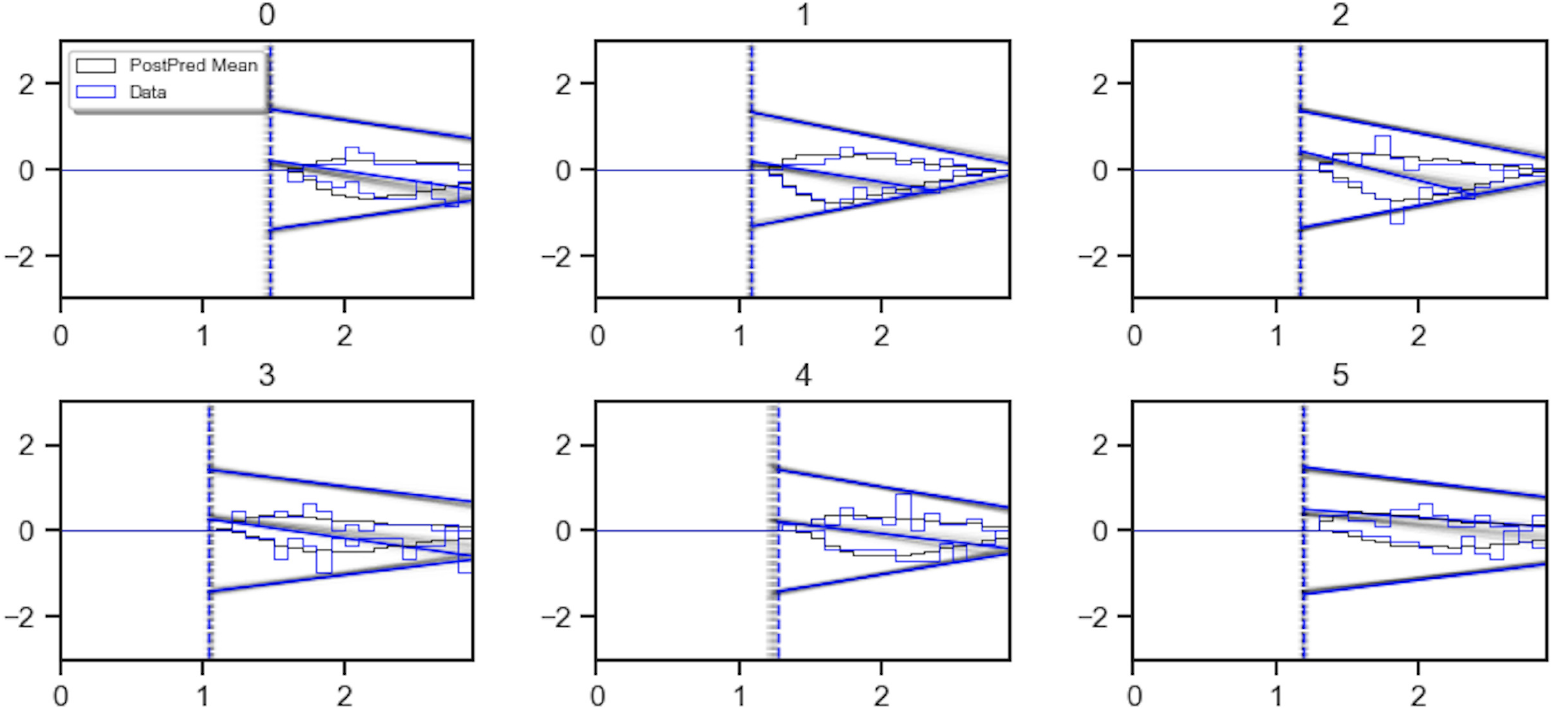
Example of a `model_plot`. This plot shows the underlying data in blue, choices and reaction times presented as a histogram (positive y-axis for choice option 1, negative y-axis or choice option 0 or −1). The black histograms show the reaction times and choices under the parameters corresponding to the posterior mean. In addition the plot shows a graphical depiction of the model corresponding to parameters drawn from the posterior distribution in black, as well as such a depiction for the *ground truth* parameters in blue, in case these were provided (e.g., if one is performing recovery from simulated data). Inclusion of the ground truth parameters distinguishes the present display form Figure 7. Various options exist to add and drop elements from this plot, the provided example corresponds to what we consider the most useful settings for purposes of illustration. Note that, in the interest of space, we only illustrate the first six subjects here.

**Figure 11.**
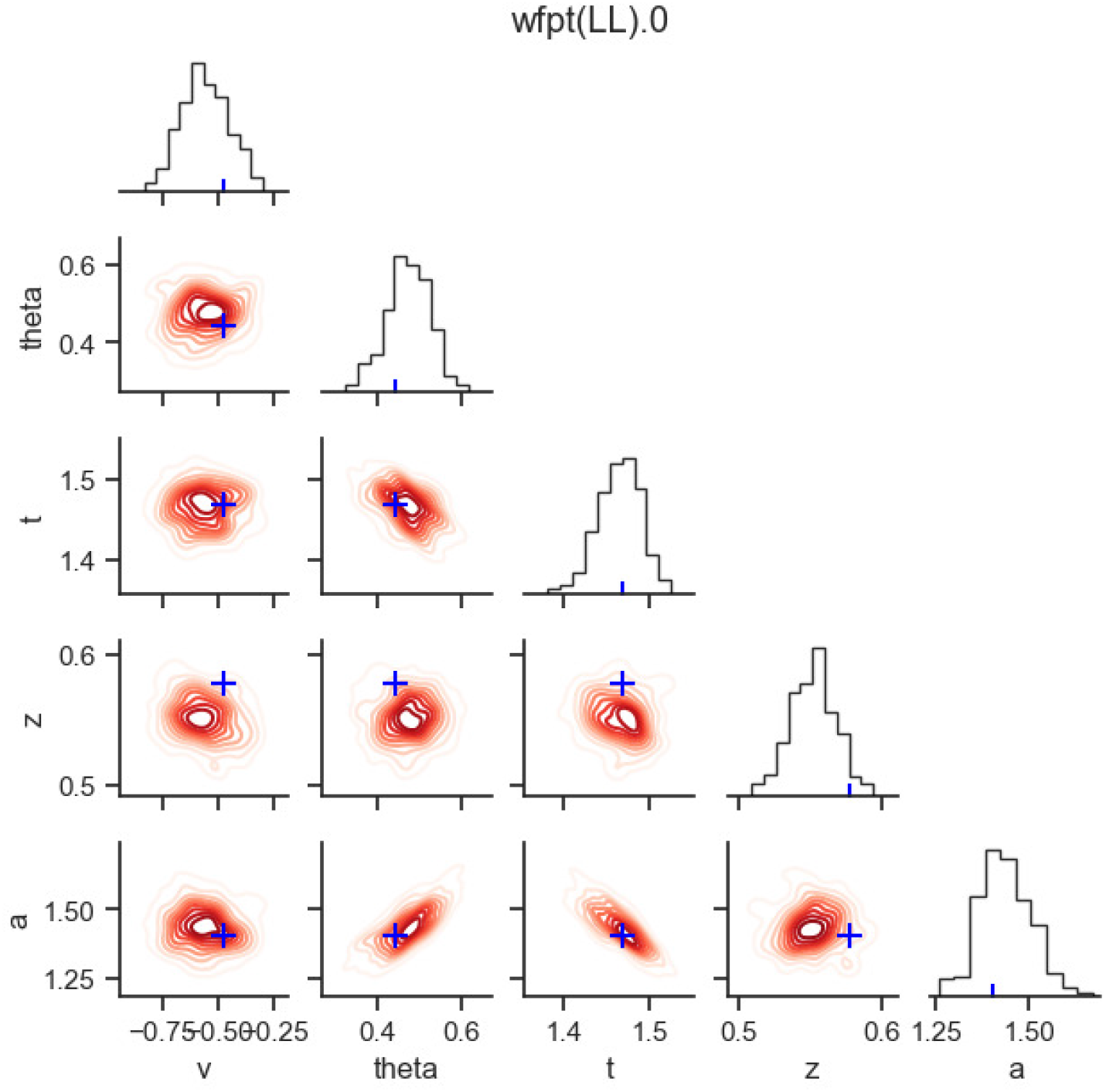
Example of a `posterior_pair_plot` in the context of parameter recovery. The plot is organized per stochastic node (here, grouped by the `‘stim’` and `‘subj_idx’` columns where in this example (`‘stim’ = ‘LL’, ‘subj_idx’ = ‘0’`). The diagonal show the *marginal posterior* of a given parameter as a histogram, adding the *ground truth* parameter as a blue tick-mark. The elements below the diagonal show pair-wise posteriors via (approximate) level curves, and add the respective *ground truths* as a blue cross.

Note how both the `plot_posterior_pair` function as well as the `plot_posterior_predictive` function take the `parameter_recovery_mode` argument to add a ground truth to the visualization automatically (the ground truth is expected to be included in the dataset attached to the HDDM model itself). The `plot_caterpillar` function needs a `ground_truth_parameter_dict` argument to add the ground truth parameters. The `simulator_h_c` function provides such a compatible dictionary of ground truth parameters. Using the set of tools in this section, we hope that HDDM conveniently facilitates application relevant parameter recovery studies.

**Codeblock 17.**
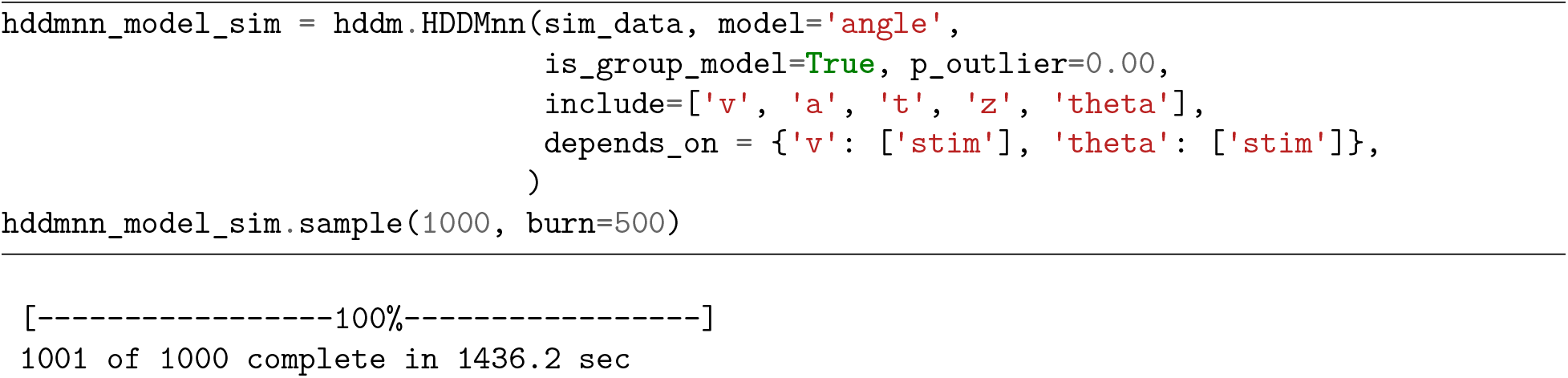
Fitting a `HDDMnn` model to synthetic data.

**Codeblock 18.**
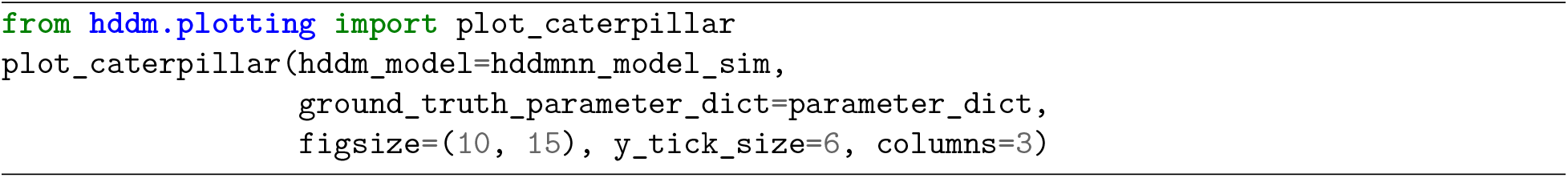
`caterpillar plot` on fit to simulated data.

**Codeblock 19.**
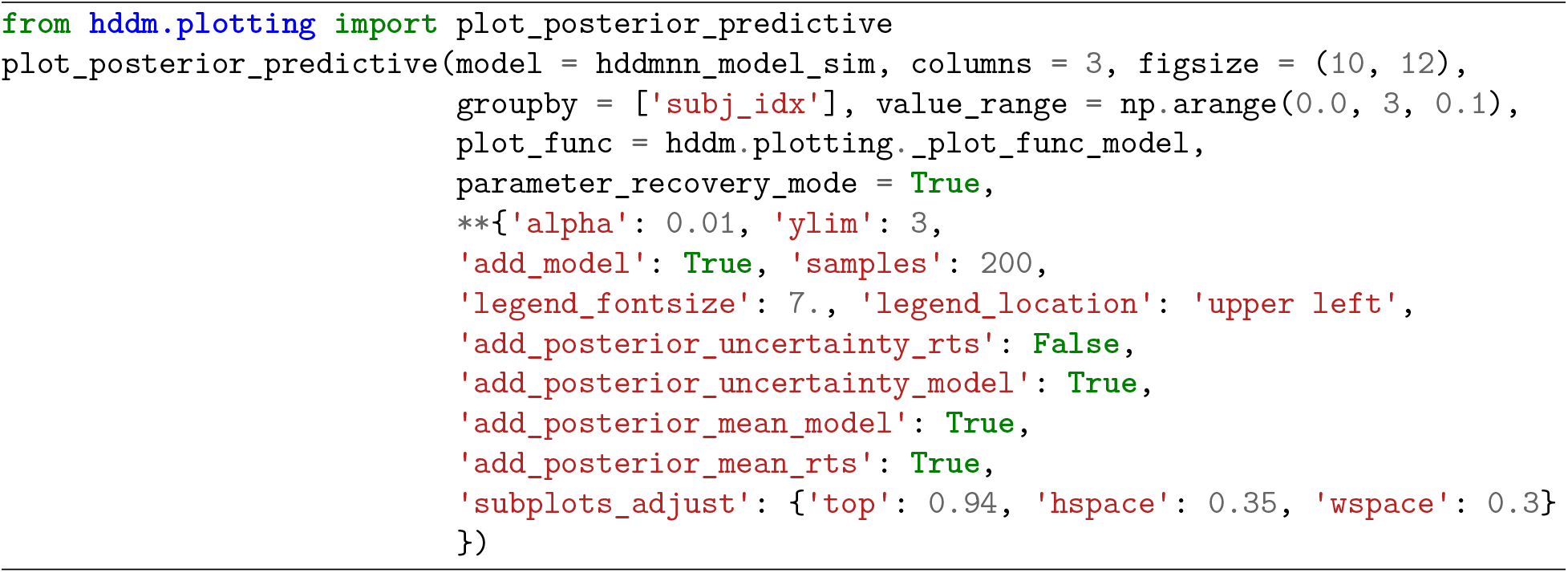
`model_plot` for fit to simulated data.

**Codeblock 20.**
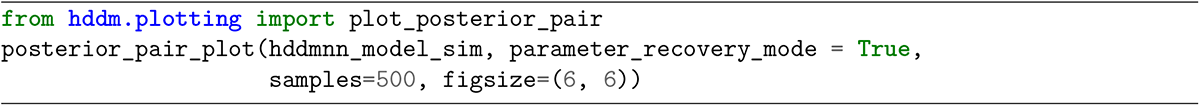
`posterior_pair_plot` for fit to simulated data.

### Adding to the bank of SSMs: User Supplied Custom Models

The new models immediately available for use with HDDM are just the beginning. HDDM allows users to define their own models via adjusting the `model_config` and the provision of custom likelihood functions. The goal of this functionality is two fold. First, we aim to make HDDM maximally flexible for advanced users, cutting down red-tape to allow creative usage. Second, we hope to motivate users to follow through with a two-step process of model integration. Step one involves easy testing of new likelihoods through HDDM, however with somewhat limited auxiliary functionality (one can generate plots based on the posterior traces, but other plots will not work because of the lack of a simulator). Step two involves sharing the model likelihood and a suitable simulator with the community to allow full integration with HDDM as well as other similar toolboxes which operate across programming languages and probabilistic programming frameworks. In future work we hope to flesh out a pipeline that allows users to follow a simple sequence of steps to full integration of their custom models with HDDM. Here, we show how to complete step one, defining a `HDDMnn` model with a custom likelihood to allow fitting a new model through HDDM. See the section on *future work* for some guidance on producing your own LAN, or contact the authors.

We start with configuring the `model_config` dictionary. We add a `“custom”` key and assign the specifics of our new model. For illustration purposes we will add the **angle** model to HDDM (even though it is already provided with the LAN extension). Additionally we need to define a basic likelihood function that takes in a vector (or matrix / `2d numpy array`) of parameters, ordered according to the list in the `“params”` key above. As an example, we load our LAN for the **angle** model (as supplied by HDDM) as if it is a custom network. Finally we can fit our newly defined *custom model*. Codeblock 21 illustrates the whole process.

**Codeblock 21.**
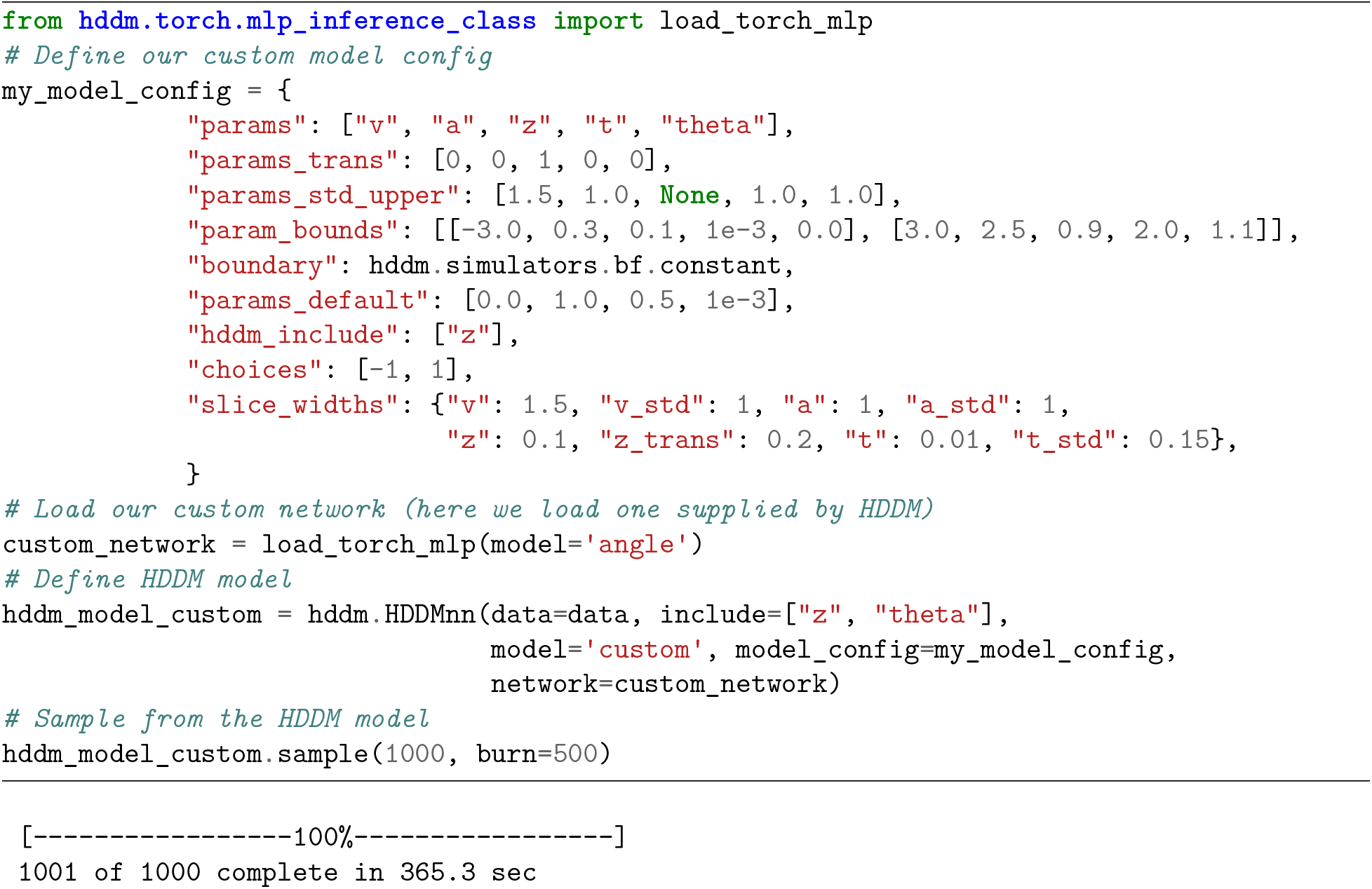
Construct and fit a HDDMnn model using a custom likelihood.

Note that the only difference to a normal call to the `hddm.HDDMnn` class is supplying appropriate model specifications for our custom likelihood. We supply the `model` argument as `“custom”`, alongside our own config dictionary to the `model_config` argument. In addition, we explicitly pass to the `network` argument, our `custom_network` defining the likelihood.

Moreover, we note that the supply of custom networks opens up multiple degrees of freedom to explore improved likelihood approximations. As an example, users may utilize LANs trained on the log-RT distributions instead of the original RT distributions of a SSM.

### Combining SSMs with Reinforcement Learning

While the previous sections focused on employing SSMs in modeling stationary environments, a host of commonly applied experimental task paradigms involve some form of learning that results from the agent’s interactions with the environment. While SSMs can be used to model the decision processes, we need additional machinery to capture the learning dynamics that arise while subjects perform such tasks. Reinforcement learning (RL) (Sutton & Barto, 2018) is one computational framework which can allow us to account for such learning processes. In reinforcement learning, researchers typically assume a simple *softmax* choice rule, informed by some ‘utility’ (or ‘goodness’) measure of taking a particular action in a given state. Mathematically, the choice probabilities are expressed as,

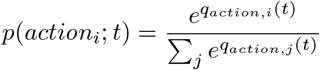

While reinforcement learning models can account for learning dynamics in basic choice behavior, the choice functions commonly employed (e.g., softmax) cannot capture the reaction time. To combine the strengths of sequential sampling models and reinforcement learning models, recent studies have used the drift diffusion model to jointly model choice and response time distributions during learning (Fontanesi et al., 2019; Pedersen & Frank, 2020; Pedersen et al., 2017). Such an approach allows researchers to study not only the across-trial dynamics of learning but also the within-trial dynamics of choice processes, using a single model. The main idea behind these models is to allow a reinforcement learning process to drive the trial-by-trial parameters of a sequential sampling model (such as the basic drift diffusion model), which in turn is used to jointly capture reaction time and choice behavior for a given trial. This can be applied in complex tasks which involve learning from feedback (see Figure 12). This results in a much more broadly applicable class of models and naturally lends itself for use in computational modeling of numerous cognitive tasks where the ’learning process’ informs the ‘decision-making process’. Indeed, a recent study showed that the joint modeling of choice and RT data can improve parameter identifiability of RL models, by providing additional information about choice dynamics (Ballard & McClure, 2019). However, to date, such models have been limited by the form of the decision model. Many RL tasks involve more than two responses, making the DDM inapplicable. Similarly, the assumption of a fixed threshold may not be valid. For example, during the early learning phase, the differences in Q-values, and hence drift rates, will be close to zero and there is little value in accumulating evidence. A standard DDM model would predict that such choices are associated with very long tail RT distributions. A more appropriate assumption would be that learners use a collapsing bound so that when no evidence is present, the decision process can terminate.

**Figure 12.**
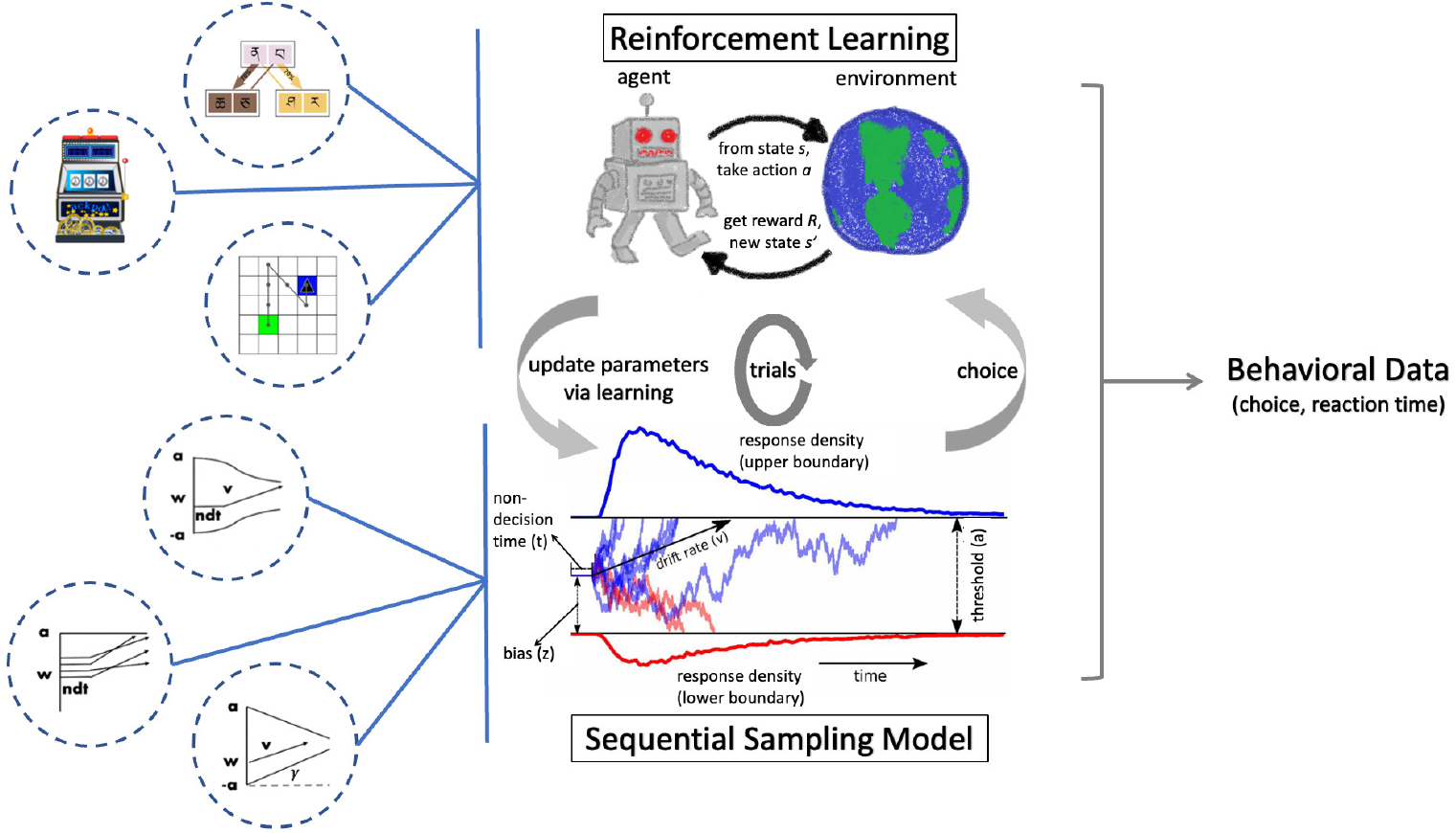
RLSSM - combining reinforcement learning and sequential sampling models.

Utilizing the power of LANs, we can further generalize the RL-DDM framework to include a much broader class of SSMs as the ‘decision-making process’. The rest of this section provides some details and code examples for these new RL-SSMs.

#### Test-bed

We test our method on a synthetic dataset of the two-armed bandit task with binary outcomes. However, our approach can be generalized to any *n*-armed bandit task given a pre-trained LAN that outputs likelihoods for the corresponding *n*-choice decision process (e.g. race models). The model employed a simple delta learning rule (Rescorla, 1972) to update the action values

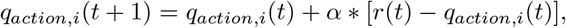

where *q_action_* (*t*) denotes expected reward (Q-value) for the chosen action at time *t*, *r*(*t*) denotes reward obtained at time *t* and *α* (referred to as *rl_alpha* in the result plots) denotes the learning rate. The trial-by-trial drift rate depends on the expected reward value learned by the RL rule. The drift rate is therefore a function of Q-value updates, and is computed by the following linking function

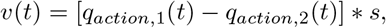

where *s* is a scaling factor of the difference in Q-values. In other words, the scalar *s* is the drift rate when the difference between the Q-values of both the actions is exactly one (Note that we refer to the scalar *s* as *v* in the corresponding figure). We show an example parameter recovery plot for this Rescorla-Wagner learning model connected to a SSM with collapsing bound in Figure 13.

**Figure 13.**
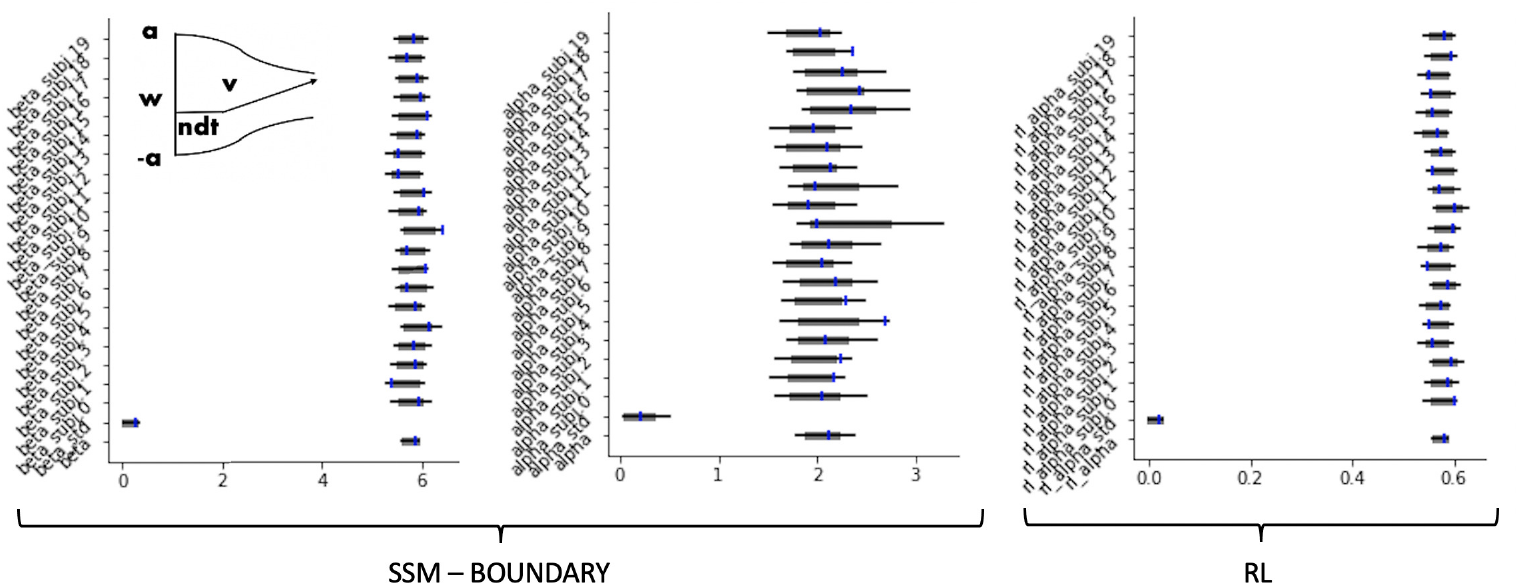
Parameter recovery on a sample synthetic dataset using RL+Weibull model. Posterior distributions for subject-level and group-level parameters are shown using caterpillar plots. The thick **black** lines correspond to 5-95 percentiles, thin black lines correspond to 1-99 percentiles. The blue tick-marks show the ground truth values of respective parameters. Note, in the interest of space, we show only a subset of the parameters of the model - the two boundary parameters **alpha** and **beta** and the reinforcement learning rate **rl_alpha**.

#### Model definitions for RL with model_config_rl

Just like the `model_config`, the new HDDM version includes `model_config_rl`, which is the central database for the RL models used in the RLSSM settings. Below is an example for simple Rescorla-Wagner updates (Rescorla, 1972), a basic reinforcement learning rule. The learning rate (referred to as ‘rl_alpha’ in Figure 13 to avoid nomenclature conflicts with the ‘alpha’ parameter in some SSMs) is the only parameter in the update rule. We do not transform this parameter (`“params_trans”` is set to 0) and specify the parameter bounds for the sampler as [0,1]. Note that for hierarchical sampling, the learning rate parameter *α* is transformed internally in the package. Therefore, the output trace for the learning rate parameter must be transformed by an inverse-logit function,

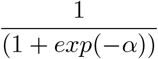

to get the learning rate values back in range [0,1]. Codeblock 22 shows us an example of such a `model_config_rl` dictionary.

**Codeblock 22.**
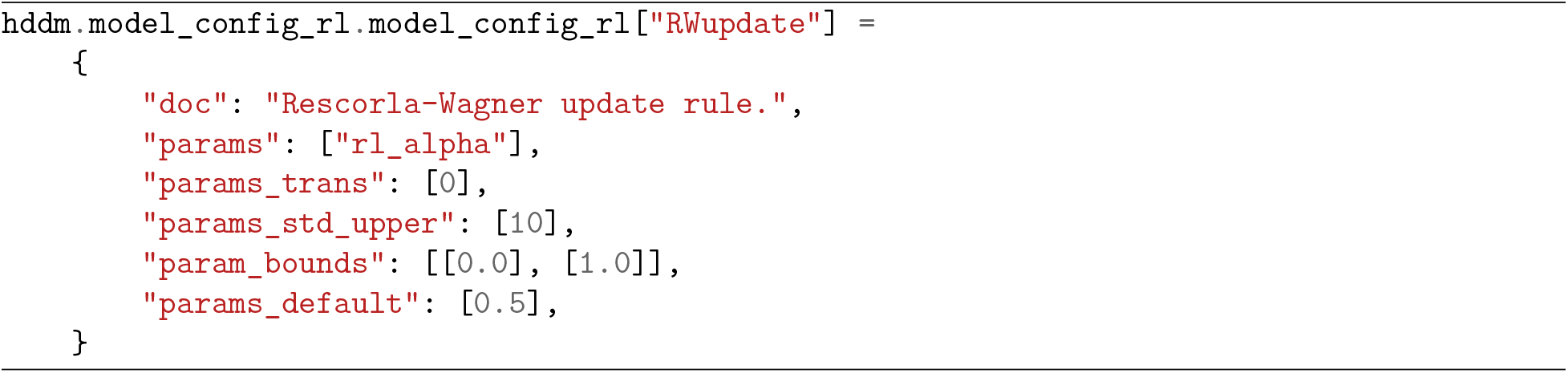
`model_config` definition for RL-SSM models.

#### Analyzing instrumental learning data

**The `HDDMnnRL` class** Running `HDDMnnRL` presents only a few slight adjustments compared to the other `HDDM` classes. First, the data-frame containing the experimental data should be properly formatted. For every subject in each condition, the trials must be sorted in ascending order to ensure proper RL updates. The column `split_by` identifies each row with a specific task condition (as integer). The `feedback` column gives the reward feedback on the current trial and `q_init` denotes the initial q-values for the model. The rest of the data columns are the same as in other `HDDM` classes. Codeblock 23 provides an example.

We can fit the data loaded in Codeblock 23 using the `HDDMnnRL` class. We showcase such a fit using the *weibull model* in conjunction with the classic Rescorla-Wagner learning rule (Rescorla, 1972). The `HDDMnnRL` class definition (shown in Codeblock 24) takes a few additional arguments compared to the `HDDMnn` class: `“rl_rule”` specifies the RL update rule to be used and `non_centered` flag denotes if the RL parameters should be re-parameterized to avoid troublesome sampling from the neck of the funnel of probability densities (Betancourt & Girolami, 2013; Papaspiliopoulos et al., 2007).

Figure 13 shows a caterpillar plot to verify the LAN-based parameter recovery on a sample RLSSM model.

#### Neural Regressors for RLSSM with the HDDMnnRLRegressor class

The new `HDDMnnRLRegressor` class, is aimed at capturing even richer (learning or choice) dynamics informed by neural activity, just like the `HDDMnnRegressor` class described above for basic SSMs. The extension works the same as the bespoke `HDDMnnRegressor` class, except that the model is now informed by a reinforcement learning process to account for the across-trial dynamics of learning. The method allows estimation of the parameters (coefficients and intercepts) linking the neural activity in a given region and time point to the RLSSM parameters.

**Codeblock 23.**
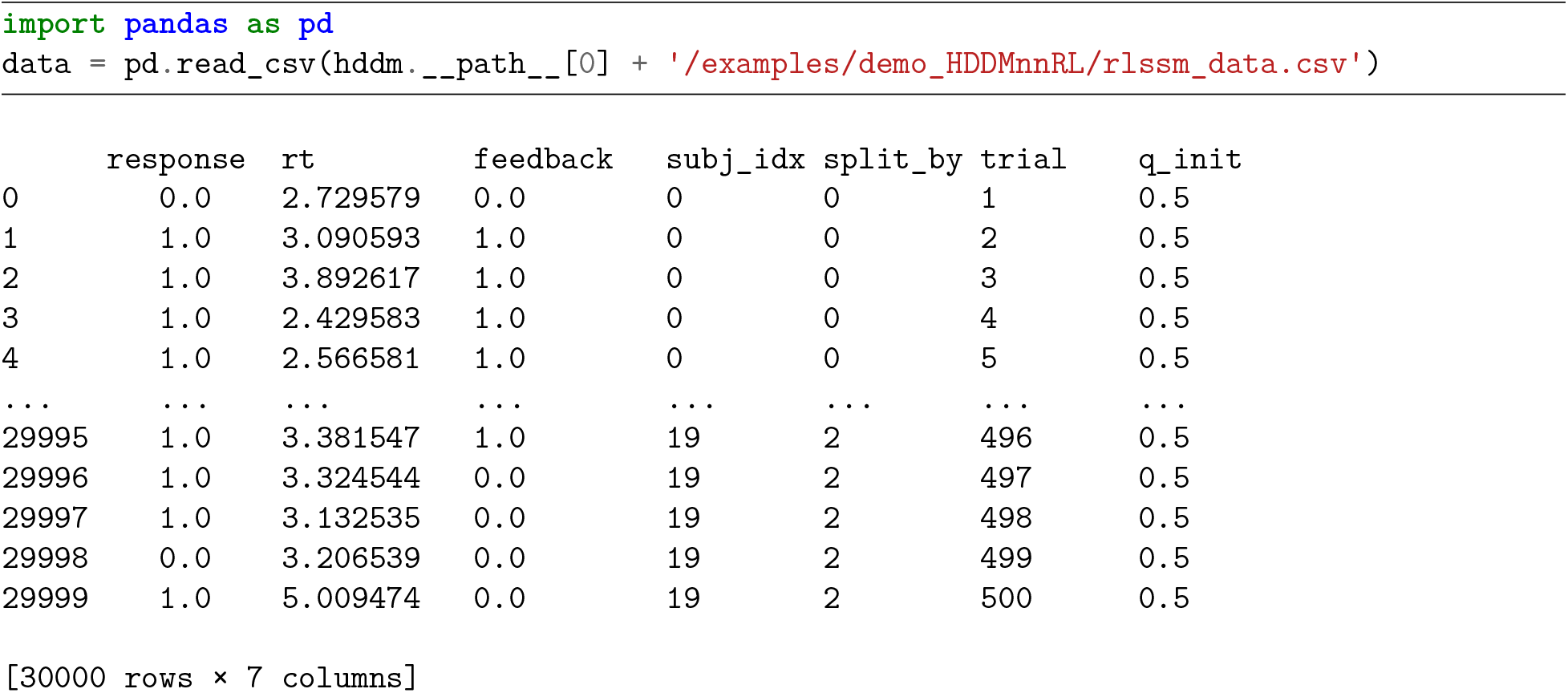
Reading in RL-SSM example data.

**Codeblock 24.**
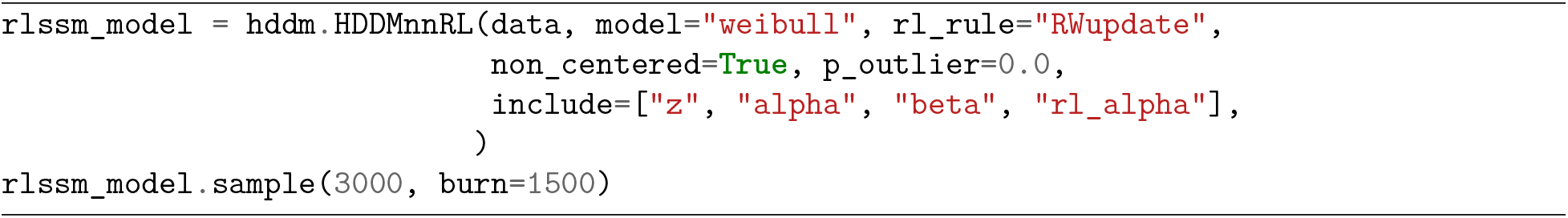
Constructing and sampling from a HDDMnnRL model.

The usage of `HDDMnnRLRegressor` class is the same as `HDDMnnRL` class except that our dataframe will now have additional column(s) for neural (or other, e.g. EEG, pupil dilation etc.) trial-by-trial covariates. Just as with the `HDDMnnRegressor` class, the model definition will also include specifying regression formulas which link covariates to model parameters. For example, if the boundary threshold parameter *a* is dependent on some neural measure *neural_reg*, Codeblock 25 shows us how to specify a corresponding `HDDMnnRLRegressor` model.

**Codeblock 25.**
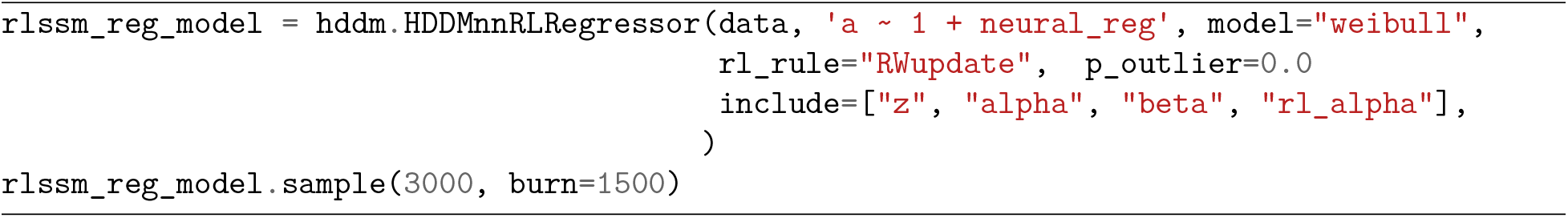
Constructing and sampling from a HDDMnnRLRegressors model.

Lastly, it is important to note that we are continually adding new functionalities to the `HDDMnnRL` and `HDDMnnRLRegressor` classes. Given the state of active development for these classes, we suggest that the users refer to the HDDM [documentation](https://hddm.readthedocs.io/en/latest/) for any updates to the usage syntax or other changes.

### More Resources

The original HDDM (Wiecki et al., 2013) paper as well as the original HDDMrl paper (Pedersen & Frank, 2020) are good resources on the basics of HDDM. The [documentation](https://hddm.readthedocs.io/en/latest/) provides examples for many complex use cases, including a [long tutorial](https://hddm.readthedocs.io/en/latest/lan_tutorial.html) specifically designed to illustrate the `HDDMnn` classes and another [tutorial](https://hddm.readthedocs.io/en/latest/demo_HDDMnnRL.html) specifically designed to showcase the `HDDMnnRL` classes. Through the [hddm user group](https://groups.google.com/g/hddm-users), an active community of HDDM users, one can find support on many problems and use cases which may not come up in the official documentation or published work.

## 5 Concluding Thoughts

We hope this tutorial can help kick-start a more widespread application of SSMs in the analysis of experimental choice and reaction time data. We consider the initial implementation with focus on LANs (Fengler et al., 2021) as a starting point, which allows a significant generalization of the model space that can be considered by experimenters. The ultimate goal however is to lead towards community engagement, providing an easy interface for the addition of custom models as a start, which could greatly expand the space of models accessible to research groups across the world. We elaborate on a few possible directions for advancements in the next section.

## 6 Limitations and Future Work

The presented extension to HDDM greatly expands the capabilities of a tried and tested Python toolbox, popular in the cognitive modeling sphere. However, using HDDM as the vehicle of choice, limitations endemic to the toolbox design remain and warrant a look ahead. First, HDDM is based on PyMC2 (Patil et al., 2010) a probabilistic modeling framework that has since been superseded by it’s successor PyMC3 (Salvatier et al., 2016) (PyMC 4.0, a rebranded PyMC has just been released too). Since PyMC2 is not an evolving toolbox, HDDM is currently bound to fairly basic MCMC algorithms, specifically a coordinate-wise slice sampler (Neal, 2003). While we have confirmed adequate posterior sampling and estimation using our LANs, estimation may be rendered more efficient if one were to leverage more recent MCMC algorithms such as Hamiltonian Monte Carlo (Hoffman & Gelman, 2014). Moreover, new libraries have emerged that act as independent functionality providers for other probabilistic programming frameworks, e.g. the ArViz (Kumar et al., 2019) python library which provides a wide array of capabilities from posterior visualizations to the computation of model comparison metrics such as the WAIC (Watanabe, 2013). Custom scripts can be used currently to deploy ArViz within HDDM. We are moreover working on a successor to HDDM (we dub it HSSM) which will be built on top of one or more of these modern probabilistic programming libraries. Second, we realize that a major bottleneck in the wider adoption of LANs (and other likelihood approximators), lies in the supply of amortizers. While our extension comes batteries included, we focused on supplying a few SSM variants of proven interest in the literature, as well as some that we used for our or lab-adjacent research. It is not HDDM, but user friendly training pipelines for amortizers, which we believe to spur the quantum leap in activity in this space. Although we are working on the supply of such a pipeline for LANs (Fengler et al., 2021), our hope is that the community will provide many alternatives. Third, we caution against uninformed use of approximate likelihoods. Before basing results of empirical studies on inference performed with LANs or other approximate likelihoods (e.g. user supplied), it is essential to test for the quality of inference that may be expected. Inference can be unreliable in manifold ways (Gelman, Rubin, et al., 1992; Geweke, 1992; Talts et al., 2018). Parameter recovery studies and calibration tests, e.g. simulation based calibration (Talts et al., 2018) should form the backbone of trust in reported analysis on empirical (experimental) datasets. To help the application of a universal standard of rigor, we are working on a set of guidelines, such as a suggested battery of tests to pass before given user supplied likelihoods should be made available to the public. Other interesting work in this sphere is emerging (Hermans et al., 2021; Lueckmann et al., 2021).

## Acknowledgments

This work was funded by NIMH grants P50 MH 119467-01 and R01 MH084840-08A1, and additionally supported by the Brainstorm Program at the Robert J. and Nancy D. Carney Institute for Brain Science. We thank Lakshmi N Govindarajan for useful tips concerning code quality. We also thank Thomas Wiecki for reviewing the additions to the HDDM package.

